# Geometric complexity and the information-theoretic comparison of functional-response models

**DOI:** 10.1101/2021.07.31.454600

**Authors:** Mark Novak, Daniel B. Stouffer

## Abstract

The assessment of relative model performance using information criteria like AIC and BIC has become routine among functional-response studies, reflecting trends in the broader ecological literature. Such information criteria allow comparison across diverse models because they penalize each model’s fit by its parametric complexity — in terms of their number of free parameters — which allows simpler models to outperform similarly fitting models of higher parametric complexity. However, criteria like AIC and BIC do not consider an additional form of model complexity, referred to as geometric complexity, which relates specifically to the mathematical form of the model. Models of equivalent parametric complexity can differ in their geometric complexity and thereby in their ability to flexibly fit data. Here we use the Fisher Information Approximation to compare, explain, and contextualize how geometric complexity varies across a large compilation of single-prey functional-response models — including prey-, ratio-, and predator-dependent formulations — reflecting varying apparent degrees and forms of non-linearity. Because a model’s geometric complexity varies with the data’s underlying experimental design, we also sought to determine which designs are best at leveling the playing field among functional-response models. Our analyses illustrate (1) the large differences in geometric complexity that exist among functional-response models, (2) there is no experimental design that can minimize these differences across all models, and (3) even the qualitative nature by which some models are more or less flexible than others is reversed by changes in experimental design. Failure to appreciate model flexibility in the empirical evaluation of functional-response models may therefore lead to biased inferences for predator–prey ecology, particularly at low experimental sample sizes where its impact is strongest. We conclude by discussing the statistical and epistemological challenges that model flexibility poses for the study of functional responses as it relates to the attainment of biological truth and predictive ability.

**Contribution to Field Statement:** The use of criteria like AIC and BIC for selecting among functional-response models is now standard, well-accepted practice, just as it is in the ecological literature as a whole. The generic desire underlying the use of these criteria is to make the comparison of model performance an unbiased and equitable process by penalizing each model’s fit to data by its *parametric complexity* (relating to its number of free parameters). Here we introduce the Fisher Information Approximation to the ecological literature and use it to understand how the *geometric complexity* of models — a form of model complexity relating to a model’s functional flexibility that is not considered by criteria like AIC and BIC — varies across a large compilation of 40 different single-prey functional-response models. Our results add caution against the simplistic use and interpretation of information-theoretic model comparisons for functional-response experiments, showing just how large an effect that model flexibility can have on inferences of model performance. We therefore use our work to help clarify the challenges that ecologists studying functional responses must face in the attainment of biological truth and predictive ability.

## 1 Introduction

> *Seek simplicity and distrust it.*
>
> Alfred North Whitehead, *The Concept of Nature*, 1919.

The literature contains thousands of functional-response experiments (DeLong & Uiterwaal, 2018), each seeking to determine the relationship between a given predator’s feeding rate and its prey’s abundance. In parallel, dozens of functional-response models have been proposed (Jeschke *et al.*, 2002; Table 1), each developed to encapsulate aspects of the variation that exists among predator and prey biologies. The desire to sift through these and identify the “best” model on the basis of data is strong given the frequent sensitivity of theoretical population-dynamic predictions to model structure and parameter values (e.g., Aldebert & Stouffer, 2018; Fussmann & Blasius, 2005). Information-theoretic model comparison criteria like the Akaike Information Criterion (AIC) and the Bayesian Information Criterion (BIC) have rapidly become the preeminent tool for satisfying this desire in a principled and quantitative manner (Okuyama, 2013), mirroring their increasing ubiquity across the ecological literature as a whole (Aho *et al.*, 2014; Ellison, 2004; Johnson & Omland, 2004). Generically, criteria like AIC and BIC make the comparison of model performance an unbiased and equitable process. For standard linear regression models (and most other models), increasing model complexity by including additional free parameters will always result in a better fit to the data. Therefore, by the principle of parsimony or because such increases in fit typically come at the cost of generality beyond the focal dataset, model performance is judged by the balance of fit and complexity when other reasons to disqualify a model do not apply (Burnham & Anderson 2002; Höge *et al.* 2018; but see Evans *et al.* 2013; Coelho *et al.* 2019)

**Table 1:**
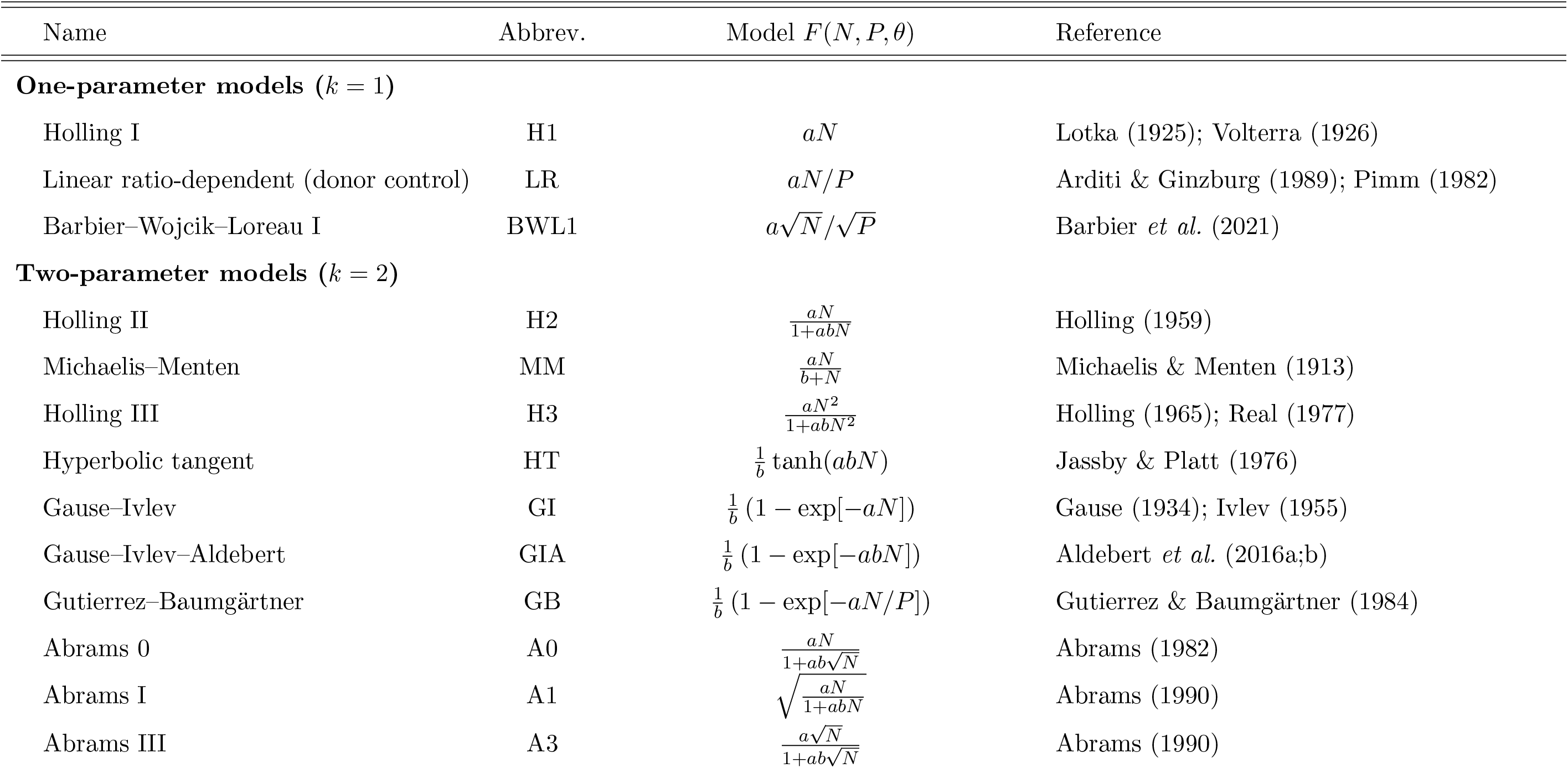
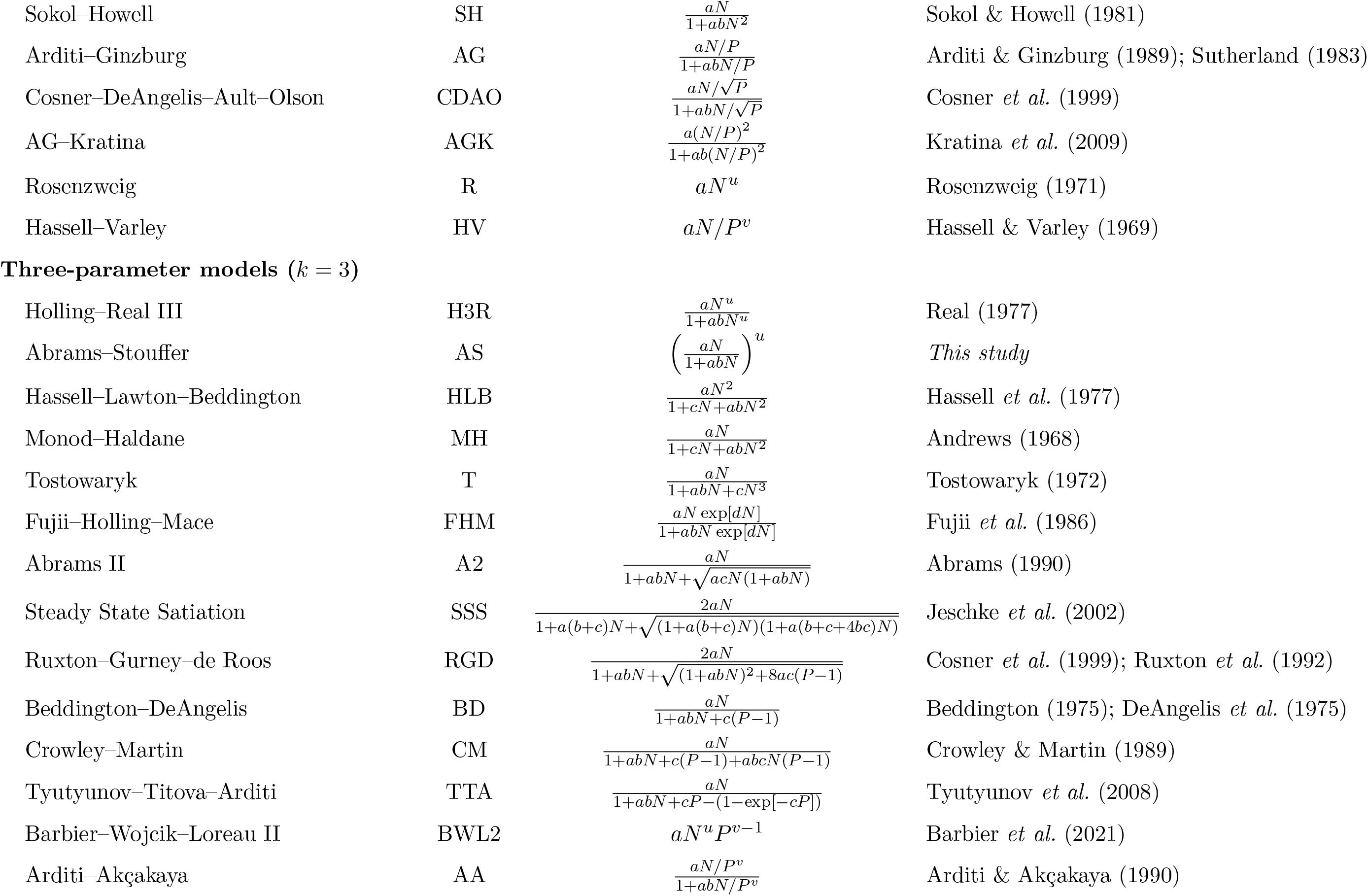
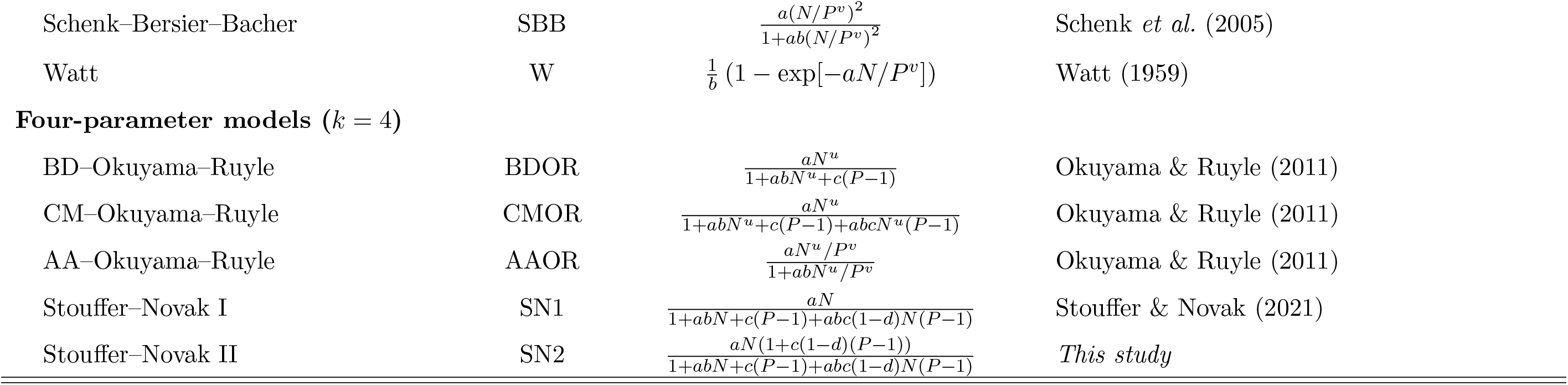
The deterministic functional-response models we considered for describing the per predator rate at which prey are eaten as a function of prey abundance *N*, predator abundance *P*, and the parameter(s) *θ*. From these per predator rates, the total count of prey eaten corresponds to the functional response, *F* (*N, P, θ*), multiplied by the number of predators *P* and the time period *T* of the experiment. The number of parameters *k* refers to the number of free parameters in each model because only these determine the mean and variance of prey eaten under the Poisson likelihood which we assumed. Note that, where appropriate, we use *P* − 1 rather than *P* for Holling-type predator-dependent models because *P* represents a count of predators in our synthetic experimental designs (rather than a density) and predator individuals cannot interfere with themselves. Original parameterizations are provided in Table S1.

| Name | Abbrev. | Model $F(N, P, \theta)$ | Reference |
| --- | --- | --- | --- |
| **One-parameter models ($k = 1$)** |  |  |  |
| Holling I | H1 | $aN$ | Lotka (1925); Volterra (1926) |
| Linear ratio-dependent (donor control) | LR | $aN/P$ | Arditi & Ginzburg (1989); Pimm (1982) |
| Barbier–Wojcik–Loreau I | BWL1 | $a\sqrt{N}/\sqrt{P}$ | Barbier *et al.* (2021) |
| **Two-parameter models ($k = 2$)** |  |  |  |
| Holling II | H2 | $\frac{aN}{1+abN}$ | Holling (1959) |
| Michaelis–Menten | MM | $\frac{aN}{b+N}$ | Michaelis & Menten (1913) |
| Holling III | H3 | $\frac{aN^2}{1+abN^2}$ | Holling (1965); Real (1977) |
| Hyperbolic tangent | HT | $\frac{1}{b} \tanh(abN)$ | Jassby & Platt (1976) |
| Gause–Ivlev | GI | $\frac{1}{b} (1 - \exp[-aN])$ | Gause (1934); Ivlev (1955) |
| Gause–Ivlev–Aldebert | GIA | $\frac{1}{b} (1 - \exp[-abN])$ | Aldebert *et al.* (2016a;b) |
| Gutierrez–Baumgärtner | GB | $\frac{1}{b} (1 - \exp[-aN/P])$ | Gutierrez & Baumgärtner (1984) |
| Abrams 0 | A0 | $\frac{aN}{1+ab\sqrt{N}}$ | Abrams (1982) |
| Abrams I | A1 | $\sqrt{\frac{aN}{1+abN}}$ | Abrams (1990) |
| Abrams III | A3 | $\frac{a\sqrt{N}}{1+ab\sqrt{N}}$ | Abrams (1990) |
| Sokol–Howell | SH | $\frac{aN}{1+abN^2}$ | Sokol & Howell (1981) |
| Arditi–Ginzburg | AG | $\frac{aN/P}{1+abN/P}$ | Arditi & Ginzburg (1989); Sutherland (1983) |
| Cosner–DeAngelis–Ault–Olson | CDAO | $\frac{aN/\sqrt{P}}{1+abN/\sqrt{P}}$ | Cosner *et al.* (1999) |
| AG–Kratina | AGK | $\frac{a(N/P)^2}{1+ab(N/P)^2}$ | Kratina *et al.* (2009) |
| Rosenzweig | R | $aN^u$ | Rosenzweig (1971) |
| Hassell–Varley | HV | $aN/P^v$ | Hassell & Varley (1969) |
| **Three-parameter models ($k = 3$)** |  |  |  |
| Holling–Real III | H3R | $\frac{aN^u}{1+abN^u}$ | Real (1977) |
| Abrams–Stouffer | AS | $\left(\frac{aN}{1+abN}\right)^u$ | *This study* |
| Hassell–Lawton–Beddington | HLB | $\frac{aN^2}{1+cN+abN^2}$ | Hassell *et al.* (1977) |
| Monod–Haldane | MH | $\frac{aN}{1+cN+abN^2}$ | Andrews (1968) |
| Tostowaryk | T | $\frac{aN}{1+abN+cN^3}$ | Tostowaryk (1972) |
| Fujii–Holling–Mace | FHM | $\frac{aN \exp[dN]}{1+abN \exp[dN]}$ | Fujii *et al.* (1986) |
| Abrams II | A2 | $\frac{aN}{1+abN+\sqrt{acN(1+abN)}}$ | Abrams (1990) |
| Steady State Satiation | SSS | $\frac{2aN}{1+a(b+c)N+\sqrt{(1+a(b+c)N)(1+a(b+c+4bc)N)}}$ | Jeschke *et al.* (2002) |
| Ruxton–Gurney–de Roos | RGD | $\frac{2aN}{1+abN+\sqrt{(1+abN)^2+8ac(P-1)}}$ | Cosner *et al.* (1999); Ruxton *et al.* (1992) |
| Beddington–DeAngelis | BD | $\frac{aN}{1+abN+c(P-1)}$ | Beddington (1975); DeAngelis *et al.* (1975) |
| Crowley–Martin | CM | $\frac{aN}{1+abN+c(P-1)+abcN(P-1)}$ | Crowley & Martin (1989) |
| Tyutyunov–Titova–Arditi | TTA | $\frac{aN}{1+abN+cP-(1-\exp[-cP])}$ | Tyutyunov *et al.* (2008) |
| Barbier–Wojcik–Loreau II | BWL2 | $aN^u P^{v-1}$ | Barbier *et al.* (2021) |
| Arditi–Akçakaya | AA | $\frac{aN/P^v}{1+abN/P^v}$ | Arditi & Akçakaya (1990) |
| Schenk–Bersier–Bacher | SBB | $\frac{a(N/P^v)^2}{1+ab(N/P^v)^2}$ | Schenk *et al.* (2005) |
| Watt | W | $\frac{1}{b} (1 - \exp[-aN/P^v])$ | Watt (1959) |
| **Four-parameter models ($k = 4$)** |  |  |  |
| BD–Okuyama–Ruyle | BDOR | $\frac{aN^u}{1+abN^u+c(P-1)}$ | Okuyama & Ruyle (2011) |
| CM–Okuyama–Ruyle | CMOR | $\frac{aN^u}{1+abN^u+c(P-1)+abcN^u(P-1)}$ | Okuyama & Ruyle (2011) |
| AA–Okuyama–Ruyle | AAOR | $\frac{aN^u/P^v}{1+abN^u/P^v}$ | Okuyama & Ruyle (2011) |
| Stouffer–Novak I | SN1 | $\frac{aN}{1+abN+c(P-1)+abc(1-d)N(P-1)}$ | Stouffer & Novak (2021) |
| Stouffer–Novak II | SN2 | $\frac{aN(1+c(1-d)(P-1))}{1+abN+c(P-1)+abc(1-d)N(P-1)}$ | *This study* |

While differing fundamentally in their underlying philosophies, motivations, and assumptions (Aho *et al.*, 2014; Höge *et al.*, 2018), both AIC and BIC implement the balance of fit and complexity in a formal manner by penalizing a model’s likelihood with a cost that depends on its number of free parameters. Specifically, for each model in the considered set of models,

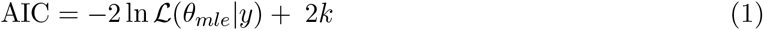

and

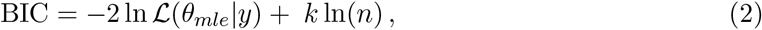

with the model evidencing the minimum value of one or the other criterion being judged as the best-performing model. For both criteria, the first term is twice the model’s negative log-likelihood (evaluated at its maximum likelihood parameter values *θ_mle_*) given the data *y*. This term reflects the model’s goodness-of-fit to the data. The second term of each criterion is a function of the model’s number of free parameters *k*. This term reflects a model’s parametric complexity. For AIC, a model’s complexity is considered to be independent of the data while for BIC it is dependent on the dataset’s sample size *n*; that is, BIC requires each additional parameter to explain proportionally more for datasets with larger sample size. The statistical clarity of the best-performing designation is typically judged by a difference of two information units between the best- and next-best performing models (Burnham & Anderson, 2002; Kass & Raftery, 1995).

An issue for criteria like AIC and BIC is that a model’s ability to fit data is not solely a function of its parametric complexity and mechanistic fidelity to the processes responsible for generating the data. This can be problematic because all models — whether it be due to their deterministic skeleton or their stochastic shell — are phenomenological to some degree in that they can never faithfully encode all the biological mechanisms responsible for generating data (see also Connolly *et al.*, 2017; Hart *et al.*, 2018). Consequently, a given model may fit data better than all other models even when it encodes the mechanisms or processes for generating the data less faithfully.

One way in which this can happen is when models differ in their flexibility. A model’s flexibility is determined by its mathematical form and can therefore differ among models having the same parametric complexity. For example, although the models *y* = *α* + *βx* and *y* = *αx^β^* have the same number of parameters and can both fit a linear relationship, the second model has a functional form that is more flexible in that it can also accommodate nonlinearities. In fact, the second model may fit some data better than the first even if the first is responsible for generating them. The chance of this happening will vary with the design of the experiment (e.g., minimizing noise and maximizing the range of *x*) and decreases as sample size increases (i.e. as the ratio of signal to noise increases). Unfortunately, sample sizes in the functional-response literature are often not large (Novak & Stouffer, 2021), and the degree to which experimental design is important given the variation in mathematical forms that exists among functional-response models has not been addressed.

Here our goal is to better understand the contrasting flexibility of functional-response models and its impact on their ranking under the information-theoretic model-comparison approach. We quantify model flexibility by geometric complexity (a.k.a. structural complexity) as estimated by the Fisher Information Approximation (FIA; Rissanen, 1996). Doing so for an encompassing set of functional-response models across experimental designs varying in prey and predator abundances, we find that geometric complexity regularly differs substantially among models of the same parametric complexity, that differences between some models can be reversed by changes to an experiment’s design, and that no experimental design can minimize differences across all models. Although choices among alternative functional-response models should be informed by motivations beyond those encoded by quantitative or statistical measures of model performance and we do not here seek to promote the use of FIA as an alternative information criterion, our results add caution against interpreting information-theoretic functional-response model comparisons merely at face value.

## 2 Materials and Methods

### 2.1 Fisher Information Approximation

The Fisher Information Approximation is an implementation of the Minimum Description Length principle (Rissanen, 1978) which Grünwald (2000) introduced as a means for making model comparisons (see Ly *et al.*, 2017; Myung *et al.*, 2006; Pitt *et al.*, 2002; for details). The Minimum Description Length (MDL) principle considers the comparison of model performance as a comparison of how well each model can compress the information that is present in data, with the best-performing model being the one that describes the data with the shortest code length. In the extreme case of random noise, no compression is possible. FIA is asymptotically equivalent to the normalized maximum likelihood which Rissanen (1996) derived to operationalize the MDL principle, but is easier to implement (Myung *et al.*, 2006). It is computed for each model as

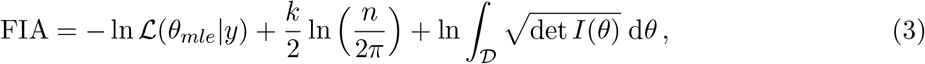

where the first term is the negative log-likelihood of the model given the data, the second term is a measure of a model’s parametric complexity that is dependent on the data via the sample size *n* (Fig. 1), and the third term is a measure of its geometric complexity (for which we henceforth use the symbol 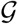). As described further in Box 1, FIA’s geometric complexity reflects a model’s ability to capture the space of potential outcomes that can be obtained given an experimental design. It thereby depends only on the model’s mathematical form and the structure underlying the observed data, but not on *n*. The contribution of geometric complexity to a model’s FIA value consequently decreases with increasing sample size relative to the contributions of the likelihood and parametric complexity. This makes the effect of geometric complexity of greatest importance for datasets with low sample sizes.

**Figure 1:**
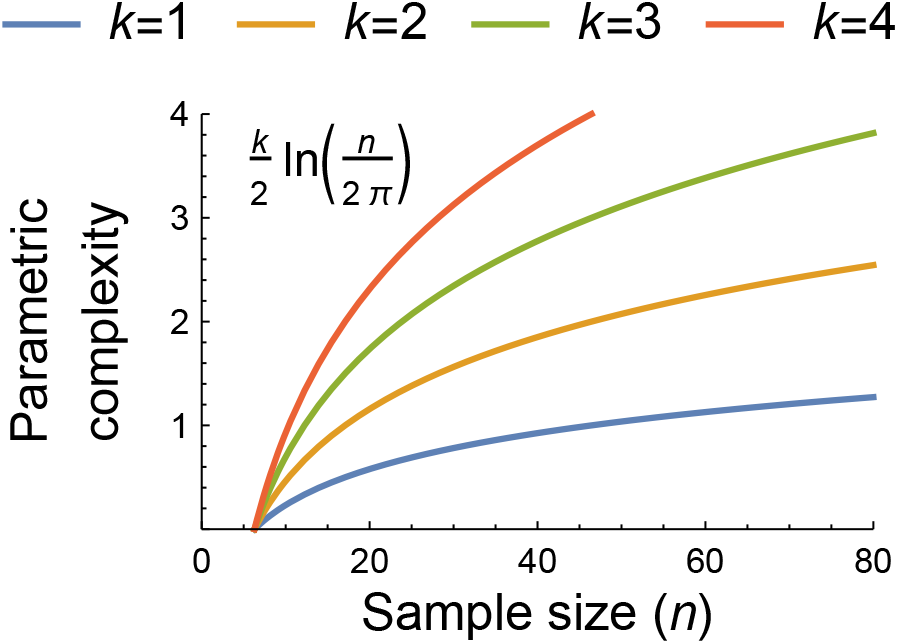
The dependence of parametric complexity on data sample size as estimated by the second term of the Fisher Information Approximation (FIA) for models with *k* = 1, 2, 3 and 4 free parameters. The potential importance of model flexibility to the information-theoretic ranking of functional-response models may be assessed by comparing their parametric and geometric complexity values or by comparing the geometric complexity values of models having the same parametric complexity because both measures of complexity are independent of the data beyond its sample size and structure (see main text and Box 1). For context, *n* = 80 was the median sample size of all functional-response datasets collated by Novak & Stouffer (2021).

For our purposes, because both parametric and geometric complexity are independent of the data beyond its sample size and experimental design, the potential importance of model flexibility to the information-theoretic ranking of models may be assessed by comparing their parametric and geometric complexity values or by comparing the geometric complexity values of models having the same parametric complexity. Because FIA converges on half the value of BIC as *n* becomes large, a one-unit difference in geometric complexity reflects a substantial impact on the relative support that two models of the same parametric complexity could receive.

##### Box 1. Unpacking the third term of the Fisher Information Approximation

As described in greater detail in Ly *et al.* (2017), Myung *et al.* (2006), and Pitt *et al.* (2002), the Fisher Information Approximation estimates the geometric complexity 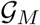 of a model *M* as the natural log of the integration (over all parameters *θ*) of the square root of the determinant of the model’s unit Fisher Information matrix *I_M_* (*θ*):

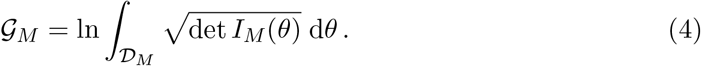

The Fisher Information matrix *I_M_* (*θ*) is a *k* × *k* matrix comprising the expected values of the second-order derivatives of the model’s negative log-likelihood function with respect to each of its *k* parameters. It therefore reflects the sensitivities of the log-likelihood’s gradient with respect to those parameters. The unit Fisher Information matrix is the expected value of these derivatives calculated across all potential experimental outcomes weighted by those outcomes’ probabilities given the parameters *θ*. When an experimental design consists of multiple treatments the expectation is averaged across these. *I_M_* (*θ*) therefore represents the expectation for a single observation (i.e. with a sample size of *n* = 1). For example, for a functional-response experiment having five prey-abundance treatment levels *N* ∈ {10, 20, 30, 40, 50} and a single predator-density level, the expectation is taken by associating a 1/5^*th*^ probability to the unit Fisher Information matrix evaluated at each treatment level. (See the *Supplementary Materials* for further details.)

The determinant of a matrix corresponds to its geometric volume. A larger determinant of the unit Fisher Information matrix therefore corresponds to a more flexible model that has higher gradient sensitivities for more of its parameters. Parameters that share all their information — such as parameters that only appear in a model as a product — result in matrix determinants of zero volume. Such non-identifiable models with statistically-redundant parameters require re-parameterization. Models can also be non-identifiable because of experimental design, such as when there is insufficient variation in predictor variables. For example, all predator-dependent functional-response models will be non-identifiable for designs entailing only a single predator abundance level (see Fig. S1).

The domain 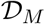 of the integral reflects the range of values that the model’s parameters could potentially exhibit. When a model is not over-specified, each location in parameter space also corresponds to a unique set of predicted model outcomes. As such, the domain of the integral reflects the space (volume) of potential experimental outcomes over which geometric complexity is calculated. Three closely related issues are pertinent in this regard:

First, a closed-form solution of the indefinite integral in Eq. (4) may not exist, and when it does it is often divergent. This means that numerical integration methods are necessary and that parameter ranges must typically be bounded (i.e. the domain 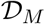 must be finite and some outcomes must be rendered “impossible”). However, how to specify bounds on mathematical grounds is not always obvious. For example, for the ratio- and consumer-dependent models such as the Hassell–Varley (HV) model, the interference strength parameter is not mathematically limited but rather can take on any non-negative value to infinity if the “attack rate” parameter is similarly unconstrained.

Second, for some experimental designs the range of parameter values may be more empirically restricted than is mathematically or even biologically permissible. For example, the handling time of the Holling Type II (H2) model (and all other models) is mathematically constrained only to be non-negative, and yet too large a handling time would mean that no prey are ever expected to be eaten except for prohibitively long experimental durations, an outcome few experimentalists would consider useful. Similarly, too large an attack rate would prevent an experimentalist from differentiating among models without the use of potentially intractable decreases in an experiment’s duration. Experimental design thereby reduces the space of possible outcomes, particularly for designs in which eaten prey are continually replaced.

Third, because a model’s geometric complexity reflects the range of parameter values which are considered possible, two models can exhibit different relative geometric complexities for different experimental designs. However, different parameterizations of the same functional form must have the same geometric complexity for a given experimental design when the permissible range of their parameters is limited equivalently (see Box 2). This is an issue because recognizing that two models simply reflect alternative parameterizations is not always easy (e.g., contrast the original formulation of the Steady State Satiation model by Jeschke *et al.* (2002) in Table S1 to our reformulation in Table 1).

In our analyses, we overcome these three issues by imposing parameter constraints in a manner that is indirect and equitable across all models. We do so by imposing the same minimum and maximum constraints on the expected number of prey eaten (thus limiting the space of potential experimental outcomes) for all models, rather than on each model’s parameters individually (see *Methods: Parameter constraints*).

##### Box 2. Imposing equitable integration limits

Different parameterizations of the same functional form should always have the same geometric complexity for a given experimental design. However, this will only be true when the range of their parameter values over which the integration of Eq. (4) is performed is limited equivalently, which can be challenging. This issue is irrelevant when solutions may be obtained in closed-form, but is not irrelevant when this is not possible, as we suspect is the case for almost all functional-response models applicable to experiments in which eaten prey are continually replaced.

The challenge of determining equitable integration limits is well-demonstrated by a comparison of the Holling and Michaelis–Menten Type II functional-response models (Fig. 2). These are typically written as

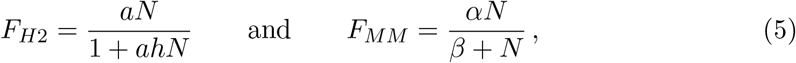

the equivalence of which is demonstrated by substituting *α* = 1/*h* (the maximum feeding rate equals the inverse of the handling time) and *β* = 1/(*ah*) (the abundance at which half-saturation occurs is the inverse of the product of the attack rate and handling time).

**Figure 2:**
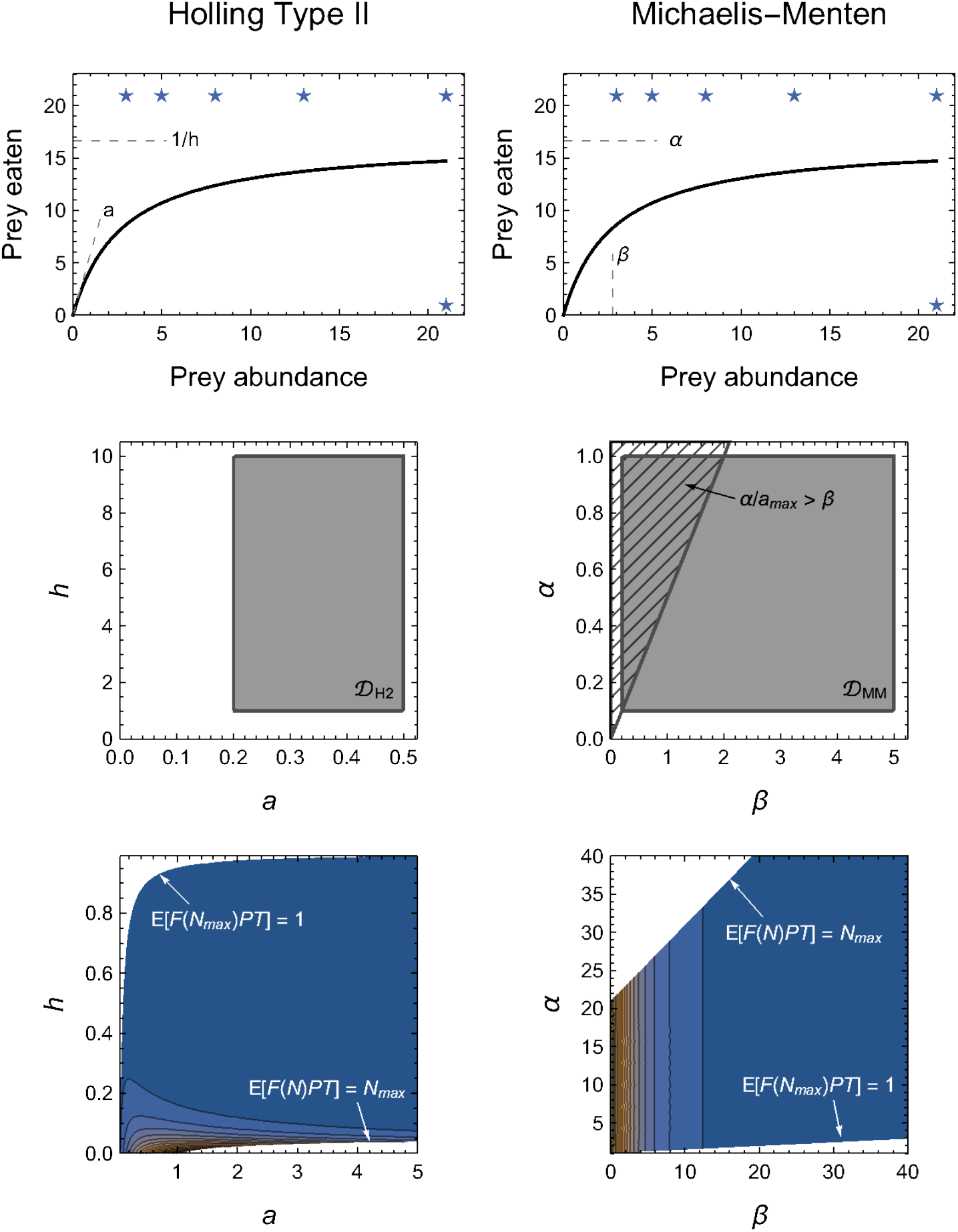
Alternative parameterizations of the same functional form should have the same geometric complexity for any given experimental design, but this will only be true in practice when their parameter domains 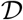 are equivalently constrained (see Box 2 for details). *Top row:* Illustration of the functional equivalence and parameter interpretations of the Holling (*left column*) and Michaelis–Menten (*right column*) models. *Middle row:* Direct constraints on 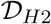 and 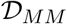 necessitate more than potentially arbitrary minimum and/or maximum limits, but must also account for the confounded relationships among parameters. *Bottom row:* We circumvent this challenge by imposing parameter constraints indirectly via the expected number of eaten prey, 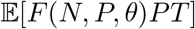. Stars in the top row indicate these limits imposed on the assumed experimental design. Colour-scale in bottom row reflects 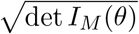 from dark blue (low values) to orange (high values), but is re-sc1a4led within each graph to visualize their contours and thus cannot be compared quantitatively.

By definition, all four parameters (*a*, *h*, *α* and *β*) are limited only in that they must be non-negative; they could each, in principle, be infinitely large (i.e. 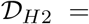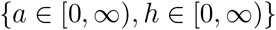 and 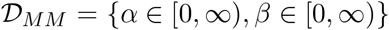). If the integral in Eq. (4) could then be computed analytically for the two models, we would always obtain 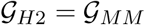 for any given experimental design.

However, because the integrals in Eq. (4) for the two models are divergent, finite limits to 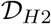 and 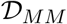 must be applied. At first glance, it may seem intuitive to impose these limits on the maximum parameter values. For example, we might consider imposing *a* ∈ [0, *a_max_*] and *h* ∈ [0, *h_max_*]. Because of their inverse relationships, doing so means that the equivalent limits for the Michaelis–Menten model are *α* ∈ [1/*h_max_,* ∞] and *β* ∈ [1/(*a_max_h_max_*), ∞], which are not finite and hence cannot solve our problem. Naively, we might therefore instead consider imposing both minima and maxima, *a* ∈ [*a_min_, a_max_*] and *h* ∈ [*h_min_, h_max_*], so that *α* ∈ [1/*h_max_,* 1/*h_min_*] and *β* ∈ [1/(*a_max_h_max_*), 1/(*a_min_h_min_*)]. This, however, does not solve a further problem in that the limits for *β* depend on the value of *α* (i.e. 1*/h*). That is, we must also impose the additional constraint that *β* > *α/a_max_* (Fig. 2), for only then will the computed 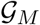 of the two models be equal.

Problems such as these only compound for models entailing a greater number of parameters. As alluded to in Box 1, our approach to circumventing these model-specific issues is to impose constraints on the expected number of eaten prey (Fig. 2), rather than on the model parameters directly (see *Methods: Parameter constraints*). That is, we require that the minimum expected number of eaten prey is no less than one prey individual in the maximum prey abundance treatment(s) (i.e. 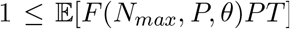 for all *P* in the experimental design) and that the maximum expected number of eaten prey is no greater than *N_max_* in any of the treatments (i.e. 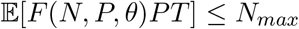 for all *N* × *P* combinations in the experimental design). Because of the mapping between parameter space and predicted model outcomes, these constraints impose natural limits for most (combinations of) parameters (e.g., the handling time or saturation parameters of all models). For other parameters, it does not impose hard limits, but nonetheless results in their contribution to 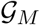 tending asymptotically to zero as their value increases (Fig. 2). This is most notably true for the “attack rate” parameter of all models.

### 2.2 Experimental designs

We computed the geometric complexity of 40 different functional-response models across a range of experimental designs. We first describe the experimental designs we considered because aspects of these also determined our manner for equitably bounding the permissible parameter space of all functional-response models (Boxes 1 & 2).

The experimental designs we considered exhibited treatment variation in prey *N* and predator *P* abundances. All designs had at least five prey-abundance levels, a minimum prey-abundance treatment of three prey individuals, and a minimum predator-abundance treatment of one predator individual. The designs varied by their maximum prey and predator abundances (*N_max_* and *P_max_*) which we achieved by correspondingly varying the number of prey and predator treatment levels (*L_N_* and *L_P_*); that is, by including higher abundance levels to smaller experimental designs. We specified the spacing between prey and predator abundance levels to follow logarithmic series. This follows the recommendation of Uszko *et al.* (2020) whose simulations showed that a logarithmic spacing of prey abundance levels performed well for the purpose of parameter estimation. We used the golden ratio (*ϕ* = 1.618 …) as the logarithmic base and rounded to the nearest integer to generate logistically-feasible abundance series that increase more slowly than typically used bases (e.g., log_2_ or log_10_). We thereby approximated the Fibonacci series (1, 1, 2, 3, 5, 8, …) on which *ϕ^n^* converges for large *n*. We varied *L_N_* between 5 and 10 levels and varied *L_P_* between 1 and 5 levels, thereby affecting *N_max_* and *P_max_* abundances of up to 233 prey and up to 8 predator individuals. We assumed balanced designs whereby all treatments are represented equally. All resulting designs are depicted in the *Supplementary Materials*.

An important aspect of experimental design which we assumed throughout our analyses was that all eaten prey are continually replaced. The constancy of available prey allowed us to treat observations as Poisson random variates and hence use a Poisson likelihood to express each deterministic functional-response model as a statistical model. This was necessary because computing geometric complexity requires an inherently statistical perspective (see Box 1 and below).

### 2.3 Functional-response models

The functional-response models we considered ranged from having one to four free parameters (Table 1). We included prey-, ratio-, and predator-dependent models that are commonly assessed in the functional-response literature, as well as many models that have received far less attention, such as those that encapsulate emergent interference, adaptive behavior, or both handling and satiation. We did not consider models that explicitly include more variables than just the abundances of a focal predator-prey pair. Given that our statistical framework was based on experimental designs within which eaten prey are continually replaced, we also did not include any models which explicitly account for prey depletion or reflect the selection of hosts by non-discriminatory parasitoids (e.g., Rogers, 1972). All but two of the considered models are previously published. The exceptions were a three-parameter model (AS) which represents an illustrative generalization of the adaptive behavior A1 model of Abrams (1990), and a four-parameter predator-dependent model (SN2) that extends the Beddington–DeAngelis and Crowley–Martin models and may be interpreted as reflecting predators that cannot interfere when feeding and can partially feed when interfering (see Stouffer & Novak, 2021).

That said, we do not concern ourselves with the biological interpretation of the models as this has been discussed extensively throughout the functional-response literature. Rather, we focus on the models’ contrasting mathematical forms. Across the different models, these forms include rational, power, and exponential functions, as well as functions that are linear, sublinear, or superlinear with respect to prey or predator abundances. To highlight their similarities, we reparameterized many models to “Holling form”, noting that different parameterizations of the same functional form have the same geometric complexity for a given experimental design (Box 2). This included models that, as originally defined, had statistically-redundant parameters (e.g., the models of Abrams, 1990), were written in “Michaelis-Menten form” (e.g., Sokol & Howell, 1981), or were written with parameters affecting divisions (e.g., we replaced 1/*c* → *c*). This also included the Steady State Satiation (SSS) model of Jeschke *et al.* (2002) for which 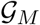 could not be computed. Fortunately the SSS model can also be derived using the citardauq formula (rather than the quadratic formula) for which 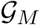 could be computed and which further reveals its similarity to the adaptive behavior A2 model of Abrams (1990) and the predator-dependent model of Ruxton *et al.* (1992). For simplicity and to further clarify similarities among models, we present all model parameters using the symbols *a, b, c*, and *d* for non-exponent parameters and *u* and *v* for exponent parameters, noting that their biological interpretations frequently differ among models.

### 2.4 Parameter constraints

As mentioned above, we assumed a Poisson statistical model in computing the geometric complexity of each deterministic functional-response model. In a context of fitting models to actual data, the consequent log-likelihood function,

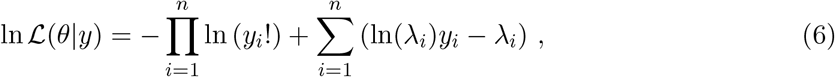

expresses the log-likelihood of a model’s parameter values given the observed data, with *λ_i_* = *F* (*N_i_, P_i_, θ*)*P_i_T*; that is, the feeding rate of a predator individual in treatment *i* (as per the focal deterministic functional-response model) times the number of predators and the time period of the experiment, which we universally set to *T* = 1. In our context of quantifying 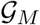, observed data is not needed because the first term of Eq. (6) drops out when taking derivatives with respect to model parameters and because *I_M_* (*θ*) involves the expected value across the space of potential experimental outcomes *y* (Box 1).

Despite this independence from data, additional information is nonetheless necessary for computing the geometric complexity of models such as those we consider here (Box 1). This information entails the range of potential outcomes that could be obtained experimentally and hence the potential parameter values that a model could exhibit (i.e. its domain 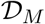 of integration). Encoding this information in an equitable manner that does not bias the inferred geometric complexity of some models over others has several potential issues associated with it (Boxes 1 and 2), particularly because the nature of our assumed experimental design (i.e. eaten prey are immediately replaced) means that the range of potential outcomes for a given model (i.e. the number of prey eaten) is theoretically infinite.

To avoid these issues, we placed no direct constraints on the parameters themselves. Rather, we specified infinite domains on the parameters (i.e. {*a, b, c, u, v*} ∈ [0, ∞) and *d* ∈ (−∞, ∞)) and instead placed constraints on them in an indirect manner by restricting the allowable outcomes predicted by the models. Specifically, we imposed the requirement that, over time-period *T*, the expected number of eaten prey in all maximum prey abundance treatments was no less than 1 (i.e. 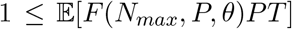 across all *P* treatments) and that the expected number of prey eaten in any treatment was no greater than the number of prey made available in the maximum prey treatment level (i.e. 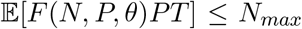 for all *N* × *P* treatment combinations). Under the assumed Poisson model, the lower bound corresponds to an expectation of observing zero prey being eaten in no greater than 37% of an experiment’s maximum prey abundance replicates (since 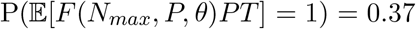). The upper bound is similarly arbitrary in a mathematical sense but seems logistically feasible since researchers are unlikely to choose a prey abundance beyond which they could not continually replace consumed individuals. For the SN1 and SN2 models, we imposed the respective additional requirement that *bd* ≤ 1/ max[*F* (*N, P, θ*)*PT*] and *b* ≤ 1/ max[*F* (*N, P, θ*)*PT*] for all treatments to maintain biologically-appropriate (non-negative) predator interference rates (Stouffer & Novak, 2021). We note that our placement of constraints on the expected number of eaten prey is similar to the use of Bayesian prior predictive checks with a joint prior distribution in that we restrict the domain of permissible parameter values based on how their conditional inter-dependencies lead to predicted model outcomes.

It is worth noting that some authors defined their models with parameters to be greater than 1, rather than 0 as we did. For example, theoreticians often assume *u* ≥ 1 for the Hill exponent of the Holling–Real Type III (H3R) model, though Real (1977) did not do so. We consider non-negative values less than one to also be biologically and statistically possible (see discussion in Stouffer & Novak, 2021). Indeed, relaxing this constraint and redefining the statistically-redundant parameters of the original A3 model (Abrams, 1990) clarifies, for example, that it is mathematically equivalent to H3R with *u* = 0.5 (even if its assumed biological mechanism differs).

### 2.5 Model comparisons

Comparisons of geometric complexity can only be made across models of the same parametric complexity; it is in conjunction with its second term that FIA enables comparisons across models in general. Therefore, for each set of models (i.e. for models with *k* = 1, 2, 3 or 4 parameters), we first assessed how an experiment’s design determined the geometric complexity of a selected “baseline” model. Because their relationships to each other and most other models are readily apparent, we chose the Holling Type I (H1) model as the baseline for the *k* = 1 models, the Holling Type II (H2) model for the *k* = 2 models, the Holling–Real Type III (H3R) and the Beddington– DeAngelis (BD) models for the *k* = 3 models (H3R for the prey-dependent models and BD for the ratio- and predator-dependent models), and the Beddington–DeAngelis–Okuyama–Ruyle (BDOR) model for the *k* = 4 models. We then compared the geometric complexity of the other models within a given set to the set’s baseline model(s) by calculating, for each experimental design, the difference between the two model’s geometric complexity values (e.g., 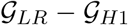). This difference enables a direct evaluation of the degree to which a model’s flexibility influences its information-theoretic ranking because it has the same units of information as the likelihood and parametric complexity terms of the FIA criterion.

### 2.6 Sensitivity to assumptions

We evaluated the sensitivity of our inferences to three aspects of experimental design, repeating our analyses for designs that

1. varied in the number of prey and predator levels (*L_N_* and *L_P_*) but kept the maximum prey and predator abundances constant at *N_max_* = 233 and *P_max_* = 5 (based on results from the main analysis);
2. used arithmetically-uniform (rather than logarithmic) series of prey and predator abundances; and that
3. relaxed the constraint on either the minimum or the maximum expected number of eaten prey by an order of magnitude (i.e. 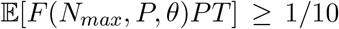 or 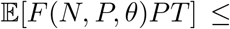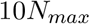).

All analyses were performed in Mathematica (Wolfram Research Inc., 2020) using the *Local Adaptive* integration method and with precision and accuracy goals set to 3 digits.

## 3 Results

### 3.1 Baseline models & equivalent models

The geometric complexity 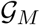 of all baseline models (H1, H2, H3R, BD and BDOR) increased with increasing *N_max_* and decreasing *P_max_* (Figs. 3–6). For these models, 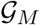 varied more greatly across the considered variation in *N_max_* than across the considered variation in *P_max_*, with at most a very weak interactive effect occurring between these. The difference in 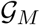 between the smallest and largest *N_max_* for a given *P_max_* varied from about 2 information units for the parametrically simplest H1 model to about 5 units for the parametrically most complex BDOR model, with the difference for the other baseline models being intermediate and roughly proportional to their number of free parameters.

**Figure 3:**
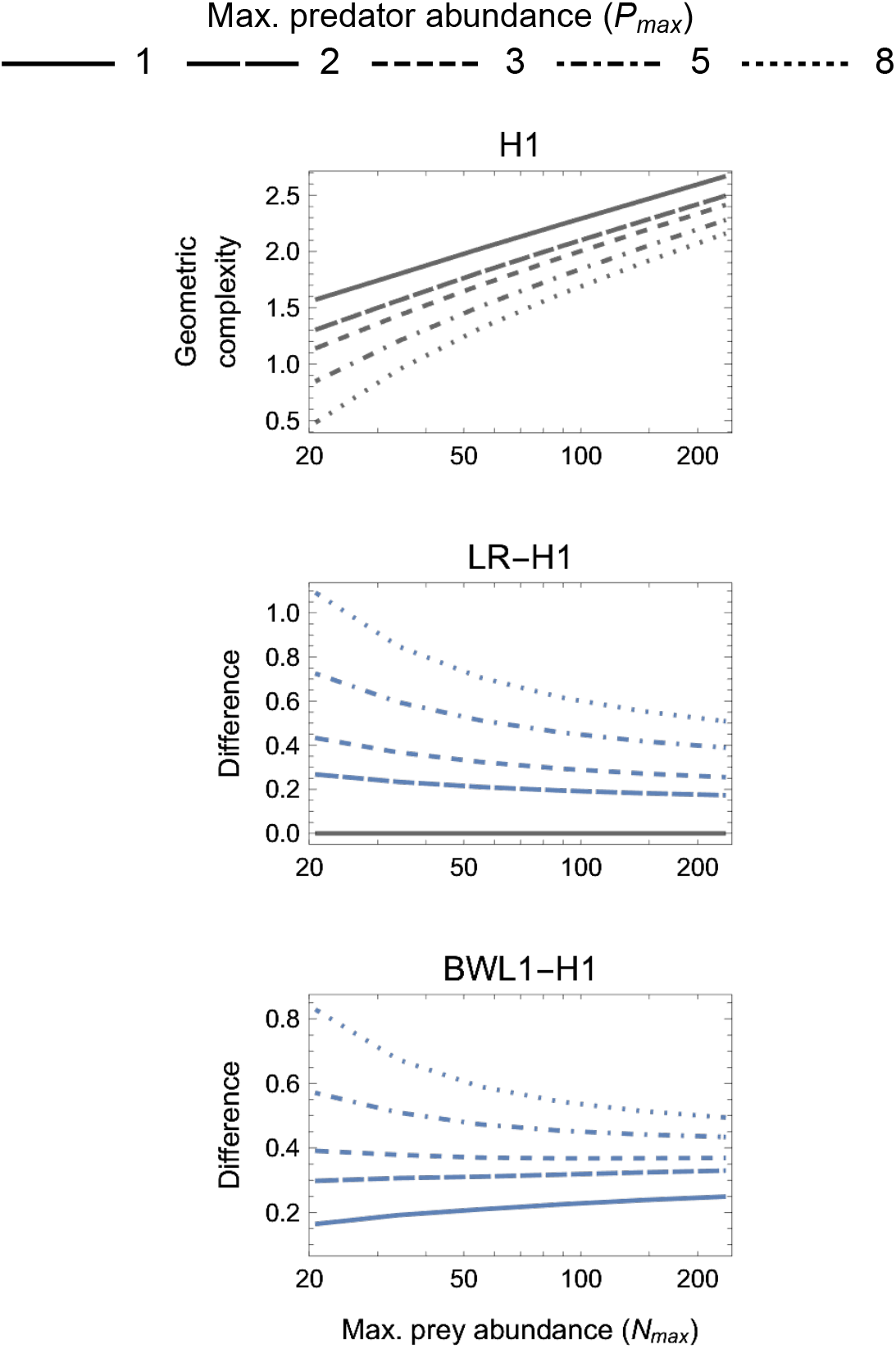
*First panel* : The geometric complexity 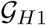 of the single-parameter (*k* = 1) base-line Holling Type I (H1) model as a function of an experiment’s maximum prey and predator abundances (*N_max_* and *P_max_*). *Other panels*: The difference in 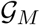 of the linear ratio-dependent (LR) model and the square-root model of Barbier *et al.* (2021; BWL1) relative to the H1 model. Positive differences reflect experimental designs for which a focal model’s mathematical flexibility would result in it being favoured by information criteria like AIC and BIC that do not consider this form of model complexity.

As expected (Box 2), alternative parameterizations of the same functional form had the same 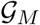 for all designs, with numerical estimation errors accounting for deviations from exact equivalence. This was demonstrated by H2 and MM as well as GI and GIA (Fig. 4), which differ only in the biological interpretation of their parameters. Likewise, all ratio-dependent models had the same 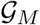 as their “corresponding” Holling-type models when there was no variation in predator abundances (e.g., 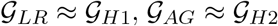 and 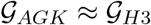 when *P_max_* = 1; Figs. 3–5).

**Figure 4:**
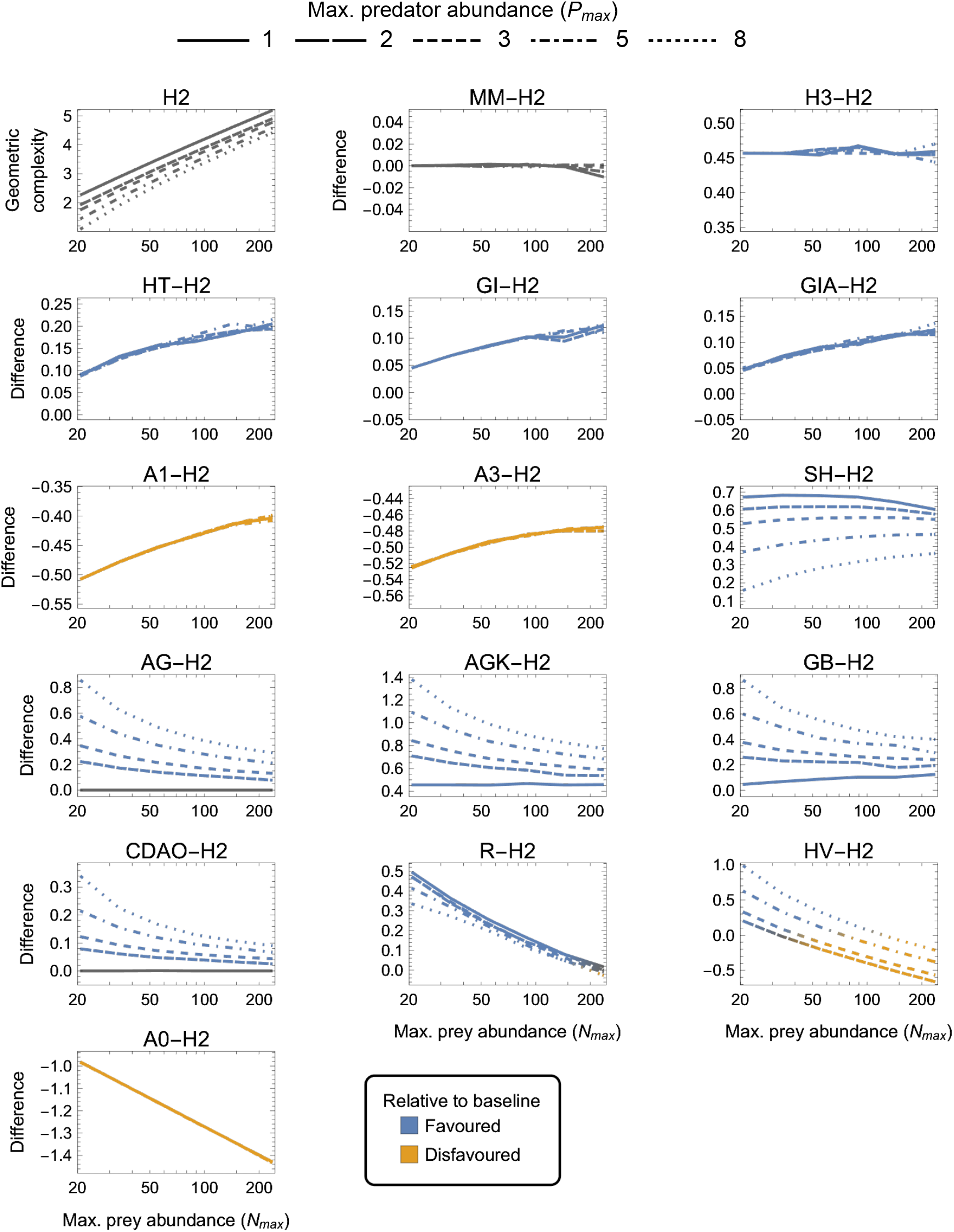
As in Fig. 3 but for two-parameter (*k* = 2) functional-response models. *First panel* : The geometric complexity 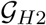 of the baseline Holling Type II model (H2) as a function of an experiment’s maximum prey and predator abundances (*N_max_* and *P_max_*). *Other panels*: The difference in 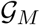 of all other two-parameter models relative to the H2 model. As a visual aid, models with greater geometric complexity than H2 are coloured in blue while those with less geometric complexity than H2 are coloured in orange.

**Figure 5:**
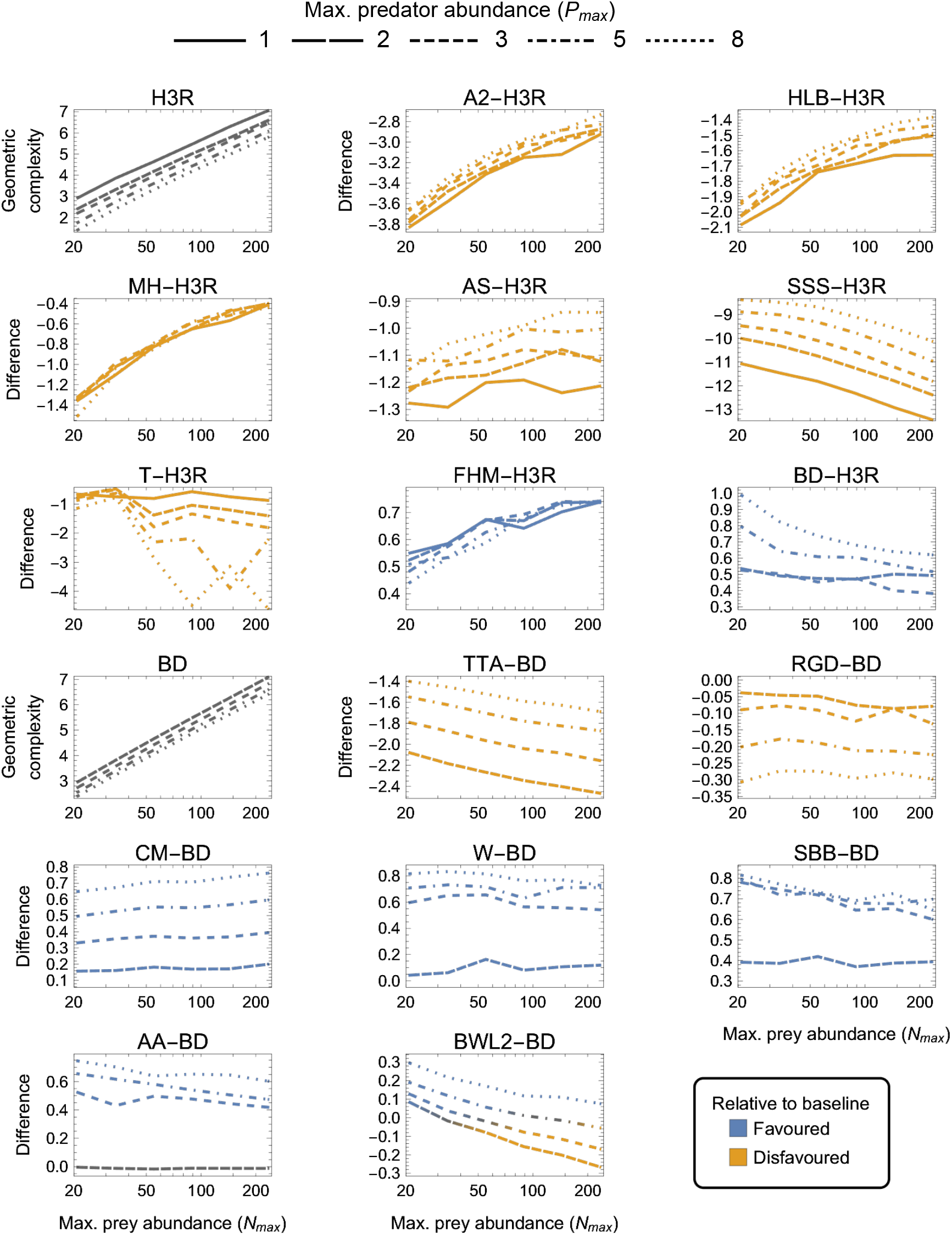
As in Fig. 3 but for three-parameter (*k* = 3) functional-response models. *First and tenth panels*: The geometric complexity 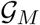 of the baseline Holling–Real Type III (H3R) and Beddington–DeAngelis (BD) models as a function of the experiment’s maximum prey and predator abundances (*N_max_* and *P_max_*). *Other panels*: The difference in 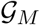 of the other three-parameter prey-dependent (top two rows) and ratio- and predator-dependent (bottom two rows) models relative to the baseline models. As a visual aid, models with greater geometric complexity than H2 are coloured in blue while those with less geometric complexity than H2 are coloured in orange.

### 3.2 One-parameter models

For the one-parameter models (Fig. 3), both 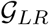 and 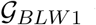 were always greater than 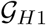 (excepting when *P_max_* = 1 for LR). The degree to which the linear ratio-dependent (LR) model was more flexible than the Holling Type I (H1) model decreased with increasing *N_max_* and decreasing *P_max_*. This was also true for the ratio-dependent BLW1 model of *Barbier et al.* (2021) when *P_max_* ≥ 3, but for *P_max_* < 3 its difference to H1 increased with increasing *N_max_*. The most equitable designs capable of differentiating among all three models therefore consisted of only two predator levels (*P_max_* = 2), entailed a 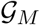 difference among models of about 0.2 information units or more, and caused LR to be slightly more flexible for small *N_max_* and BWL1 more so for large *N_max_* relative to H1. The least equitable design entailed large *P_max_* and small *N_max_* and caused the geometric complexity of LR and BWL1 to exceed that of H1 by more than 1 and 0.8 information units, respectively.

### 3.3 Two-parameter models

There were four categories of two-parameter models qualitatively distinguished by whether they exhibited equivalent, higher, lower or a design-dependent 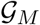 relative to the H2 baseline model (Fig. 4):

i. MM was equivalent to H2 for all designs (as already mentioned above);
ii. H3, HT, GI, GIA, SH, AG, AGK, GB, CDAO and R were more flexible than H2 for all designs (had higher 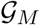, excepting for *P_max_* = 1 where 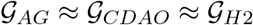);
iii. A0, A1 and A3 were less flexible than H2 for all designs (had lower 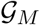); and
iv. HV was more flexible than H2 for small *N_max_* designs and less flexible for large *N_max_*, with large and small *P_max_* designs respectively increasing and decreasing its relative flexibility more greatly.

H3 was the only model for which the difference from H2 was insensitive to experimental design, always being about 0.45 information units. For HT, GI, GIA, A0, A1 and A3, the difference to H2 was insensitive to *P_max_*, but while it increased with increasing *N_max_* for HT, GI, GIA and A0 (making small *N_max_* designs the most equitable), it decreased with increasing *N_max_* for A1 and A3 (making large *N_max_* designs the most equitable). The degree to which AG, AGK, GB, CDAO and R were more flexible than H2 decreased with increasing *N_max_*, but while it increased with increasing *P_max_* for AG, AGK, GB and CDAO (making large *N_max_*, small *P_max_* designs the most equitable), it decreased — albeit weakly — with increasing *P_max_* for R. For SH, the difference to H2 first increased from small to intermediate *N_max_* then slowly decreased from intermediate to large *N_max_*, but was always minimized by large *P_max_*. Small *N_max_*, large *P_max_* designs were therefore the most equitable for SH. Finally, for HV, which was either more or less flexible than H2 depending on design, the most equitable designs spanned *N_max_* ≈ 30 for *P_max_* = 2 to *N_max_* ≈ 120 for *P_max_* = 8. Overall, A0 and AGK exhibited the greatest potential disparity in flexibility relative to H2, respectively being less and more flexible by about 1.4 information units under their least equitable design. The greatest potential disparity among all considered two-parameter models was about 2 information units and occurred between HV and A0 for small *N_max_*, large *P_max_* designs in favour of HV.

### 3.4 Three-parameter models

Noting that all predator-dependent models are non-identifiable for *P_max_* = 1 designs (Fig. S1), there were three categories of three-parameter models that were qualitatively distinguished by whether they exhibited higher, lower or a design-dependent 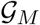 relative to the two baseline models — H3R for prey-dependent models and BD for ratio- and predator-dependent models (Fig. 5):

i. FHM and BD were more flexible than H3R, and CM, W, SBB and AA were less flexible than BD, for all designs (excepting for *P_max_* = 2 where 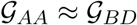);
ii. A2, HLB, MH, AS, SSS and T were less flexible than H3R, and TTA and RGD were less flexible than BD, for all designs; and
iii. BWL2 was more flexible than BD for small *N_max_*, large *P_max_* designs and was less flexible for large *N_max_*, small *P_max_* designs.

For the ratio- and predator-dependent models, differences to BD were more sensitive to variation in *P_max_* than to variation in *N_max_*. The degree to which CM, W, SBB and AA were more flexible than BD increased with increasing *P_max_*, reaching a difference in geometric complexity of 0.8 information units at *P_max_* = 8. For these models, the most equitable design therefore entailed small *P_max_* regardless of *N_max_*, but for TTA and RGD, for which the difference to BD decreased with increasing *P_max_*, it was designs entailing large *P_max_* which reduced their lower geometric complexity the least (by no less than 1.4 and up to 2.9 information units). The degree to which the prey-dependent AS, SSS and T models were less flexible than H3R was also more sensitive to variation in *P_max_* than in *N_max_*, but the degree to which A2, HLB and MH were less flexible and the degree to which FHM was more flexible was relatively insensitive to variation in *P_max_*. As *N_max_* increased, SSS and T became less flexible than H3R, A2, HLB, MH and AS became less inflexible relative to H3R, and FHM became more flexible than H3R. For BWL2, which could either be more or less flexible than H3R depending on design, the most equitable designs spanned those that had the largest considered *N_max_* when *P_max_* was large to those that had the smallest considered *N_max_* when *P_max_* was small. Overall, SSS and RGD exhibited the greatest potential disparity relative to their H3R and BD baselines, respectively differing in their geometric complexity by about 13 and almost 2.9 information units for the least equitable designs. The greatest potential disparity among all other considered three-parameter models was about 11 information units and occurred between SSS and CM for large *N_max_*, large *P_max_* designs in favour of CM.

### 3.5 Four-parameter models

Finally, among the four-parameter models, which exhibited the greatest amounts of numerical estimation noise (Fig. 6):

i. AAOR was more flexible than BDRO for all designs (had higher 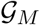);
ii. SN1 and SN2 were less flexible than BDRO for all designs (had lower 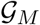); and
iii. CMOR tended to be more flexible for large *N_max_*, large *P_max_* designs and less flexible for small *N_max_*, small *P_max_* designs.

**Figure 6:**
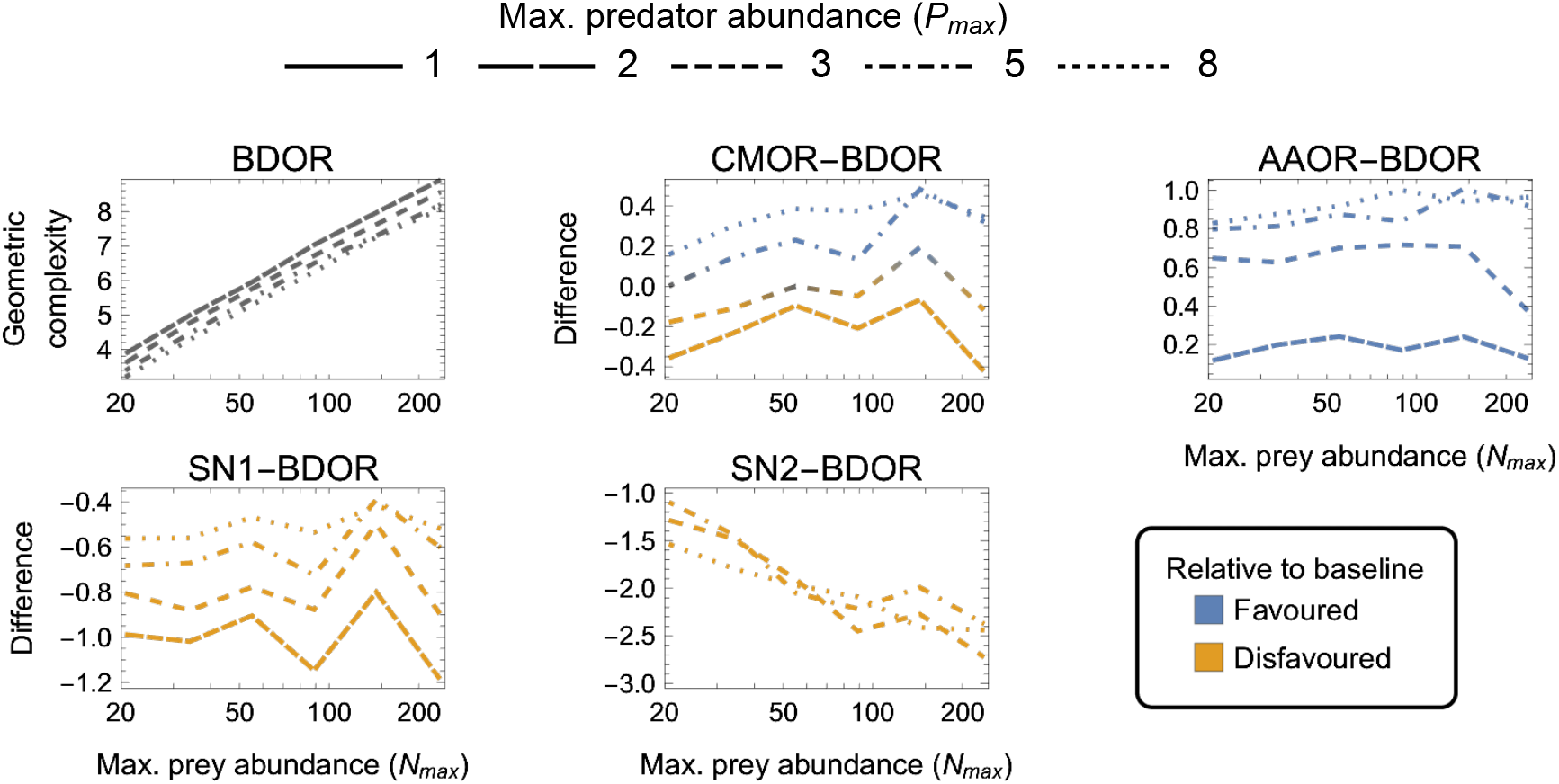
As in Fig. 3 but for four-parameter (*k* = 4) functional-response models. *First panel* : The geometric complexity 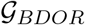 of the Beddington–DeAngelis–Okuyama–Ruyle (BDOR) model as a function of the experiment’s maximum prey and predator abundances (*N_max_* and *P_max_*). *Other panels*: The difference in 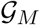 of all other four-parameter models relative to the BDOR model. As a visual aid, models with greater geometric complexity than H2 are coloured in blue while those with less geometric complexity than H2 are coloured in orange.

For CMOR, AAOR and SN1, the difference to BDOR was less sensitive to variation in *N_max_* than to variation in *P_max_*, but the opposite was true for SN2. Further, while the degree to which AAOR was more flexible than BDRO was minimized by *P_max_* = 2 designs (to about 0.2 information units), the degree to which SN1 was less flexible than BDRO was minimized by *P_max_* = 8 designs (to about 0.5 information units). SN2 was non-identifiable for designs having *P_max_* ≤ 3 (Fig. S1), but for *P_max_* > 3 designs it was less flexible by at least 1 information unit. The most equitable designs for CMOR and BDOR entailed intermediate predator abundances (*P_max_* = 3–5). Overall, the greatest potential disparity to the BDOR baseline model occurred for the SN2 model (about 2.5 information units) at the largest *N_max_*. The greatest potential disparity among all considered four-parameter models occurred for the SN2 and AAOR models (about 3.5 information units) for the largest *N_max_*, largest *P_max_* design in favour of AAOR.

### 3.6 Sensitivity analyses

Fixing *N_max_* = 233 and *P_max_* = 5 and varying the number of prey and predator treatment levels (*L_N_* and *L_P_*) to below the numbers used in our primary analysis showed that 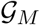 was relatively insensitive to variation in *L_N_* for most models (Figs. S2–S5). In contrast, the degree to which models were more or less flexible relative to their baseline model was far more sensitive to variation in *L_P_*. For most of the *L_P_*-sensitive models, decreasing *L_P_* increased their difference to the baseline model, but for an almost equal number the difference decreased. The largest effects of *L_P_* most often occurred when reducing from two predator levels (*P* ∈ {1, 2}) to only a single-predator level (or the corresponding reduction of three to two levels for the four-parameter models). Setting aside these last-mentioned and in some ways trivial changes to *L_P_*, the greatest effect of changing *L_N_* and *L_P_* was to change the relative geometric complexity of models and their baseline models by up to about 0.6 information units (excepting T and SSS for which changes of up to 2.5 units occurred).

The use of designs with arithmetic rather than logarithmic spacings of prey and predator abundances also had little to no effect on the geometric complexity of models relative to their baselines (Figs. S6–S9). The notable exceptions included the manner in which (i) HV was more flexible than H2 (arithmetic spacings making HV invariably more flexible rather than more or less flexible depending on *N_max_* and *P_max_*), (ii) BD was more flexible than H3R (arithmetic spacings making it more flexible for large rather than small *N_max_*), and (iii) CMOR was more flexible than BDOR (arithmetic spacings making CMOR invariably less flexible rather than more or less flexible depending on *N_max_* and *P_max_*).

Finally, relaxing the indirect constraints we imposed on the range of potential experimental outcomes (i.e. model parameters) by changing the minimum or the maximum expected number of eaten prey by an order of magnitude had similarly little effect (Figs. S10–S17). The notable consequences were that increasing the maximum expected number of eaten prey across all treatments from *N_max_* to 10 *N_max_* caused (i) CDAO to become less rather than more flexible than H2, (ii) T and W to be more or less flexible than H3R and BD in a design-dependent rather than design-independent manner; (iii) CMOR to become more flexible than BDOR for a greater range of designs, and (iv) 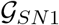 and 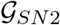 to no longer be estimable, even after a month of computation on a high-performance computing cluster.

## 4 Discussion

The functional-response literature is replete with models, even among those that only consider variation in the abundances of a single predator-prey pair (Table 1, Jeschke *et al.*, 2002). Each of these many deterministic models was proposed to encapsulate a different aspect of predatorprey biology, though frequently even very different biological processes lead to very similar or even the same model form (Table 1). Information-theoretic criteria, which balance model fit and complexity, represent the principal, most general, and most accessible means for comparing the statistical performance of these models when they are given a statistical shell and confronted with data (Okuyama, 2013). The primary contribution of our analyses is to show that existing models, independent of the biology they are meant to reflect, frequently also differ in their flexibility to fit data, even among models having the same parametric complexity. Differences in model flexibility as assessed by the geometric complexity term 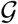 of the FIA criterion were frequently greater than 0.5 information units, spanned values up to 13 information units, and for several models were never below 1 information unit even for the most equitable of considered experimental designs. Secondarily, our analyses demonstrate just how dependent a model’s flexibility can be on the experimental design of the data (i.e. what the range and combinations of prey and predator abundances are). In some instances this design dependency was great enough to cause models that were less flexible than other models for some experimental designs to become more flexible than the same models for different designs.

Our use of the FIA criterion allows us to contextualize the importance of this variation in flexibility in two rigorous and quantitative ways: First, we can compare 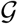 among models of the same parametric complexity for a given experimental design assuming their goodness-of-fit to a hypothetical dataset to be the same. In this scenario, the potential significance of model flexibility to the information-theoretic comparison of functional-response models is evidenced in a general manner by the fact that a 2-unit difference in AIC or BIC among competing models — equivalent to a 1-unit difference in FIA — represents “substantial” support (a weight-of-evidence of 2.7 to 1) for one model over another (Burnham & Anderson, 2002). (Such a difference reflects a probability of 0.73 that the first of only two competing models is “better” than the other.) Second, we can compare 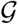 to a model’s parametric complexity for hypothetical datasets of differing sample size assuming its goodness-of-fit to these data remains the same. In this scenario, the potential significance of model flexibility to the inferences of functional-response studies performed in the past is evidenced by the fact that our estimated differences in 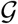 are comparable to the values of parametric complexity that are associated with the median and even maximum sample sizes seen in the large collection of datasets recently compiled by Novak & Stouffer (2021) (Table 2). That is, as feared by Novak & Stouffer (2021), sample sizes among existing empirical datasets are often sufficiently small that the likelihood and parametric complexity differences of many models is unlikely to have sufficiently out-weighed the influence of their functional flexibility in determining their information-theoretic rankings.

**Table 2:** The value of FIA’s parametric complexity term (the second term of Eq. (3) depicted in Fig. 1) for models of *k* = 1, 2, 3 and 4 parameters evaluated at the sample sizes of the smallest (*n* = 10), median (*n* = 80), and largest (*n* = 528) sized datasets in the set of 77 functional-response datasets having variation in both prey and predator abundances compiled by Novak & Stouffer (2021). These values serve as reference for gauging the magnitude differences in geometric complexity between models reported here and, thereby, for judging the likely influence of model flexibility on prior inferences of relative model performance using AIC and BIC.

| $k$ | Sample size ($n$) | | |
| --- | --- | --- | --- |
|  | 10 | 80 | 528 |
| 1 | 0.2 | 1.3 | 2.2 |
| 2 | 0.5 | 2.5 | 4.4 |
| 3 | 0.7 | 3.8 | 6.6 |
| 4 | 0.9 | 5.1 | 8.9 |

### 4.1 What makes models (in)flexible?

Given that the influence of model flexibility on information-theoretic model comparisons of the past is likely substantial, that its influence will likely not change dramatically in the future given the logistical challenges of standard experimental approaches, and because there is no experimental design that can make the comparison of functional-response models universally equitable with respect to their flexibility, an important question is: What aspects of their mathematical formulation make models more or less flexible for certain experimental designs?

For the one-parameter models the answer is relatively accessible given the specifics of our analyses. The linear ratio-dependent (LR) model is more flexible than the Holling Type I (H1) model because the division of prey abundances by a range of predator abundances allows a greater range of parameter *a* (“attack rate”) values to satisfy the condition that the resulting expected numbers of eaten prey will lie within our specified minimum and maximum bounds (i.e. satisfying both 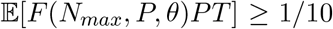 and 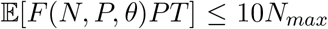). Relative to H1 for which high *N_max_* and low *P_max_* maximize the potential range of attack rates that an individual predator could express in an experiment, having many predators “interfering” in a ratio-dependent manner enables each individual predator to express an even greater attack rate without all predators in total consuming too many prey. The effects on the maximum versus the minimum prey eaten are asymmetric in magnitude (i.e. the maximum potential value of *a* increases more than the minimum potential value of *a*) because division by *P* in LR has an asymmetric effect on the per predator number of prey eaten (relative to the multiplication by *P* that is common to all models); it is symmetric only on a logarithmic scale. The magnitude of this effect is dampened in the BWL1 model of Barbier *et al.* (2021) because it entails a ratio of the square roots of (is sublinear with respect to) prey and predator abundances, making BWL1 more flexible than H1 but less flexible than LR.

The same rationale applies to all other models and explains the varied (in)sensitivities that their model flexibility has with respect to experimental design. That said, the situation is often more complicated for models with multiple parameters because of (i) the interdependent influences that parameters have on the number of prey that are eaten, and (ii) the fact that, for some models, the minimum and the maximum boundaries on the expected number of eaten prey come into play at different points in parameter- and species-abundance space.

For example, for the Holling Type II (H2) model, requiring that at least one prey on average be eaten in the highest prey abundance treatments causes high handling times to impose a lower limit on each individual’s attack rates only if and when prey abundances are sufficiently high to affect saturation. The Holling Type III (H3) model experiences this same effect as well, hence its *relative* flexibility is insensitive to variation in maximum prey abundances. H3 is nonetheless more flexible than H2 because it is superlinear with respect to prey abundance (when handling times or prey abundances are low) and can therefore satisfy the minimum of one-prey-eaten-per-predator constraint for smaller attack rate values than can H2. Similarly, the exponential form of the Gause–Ivlev models (GI and GIA) makes them more flexible than H2 because they are superlinear with respect to prey abundance, while the A1 and A3 models of Abrams (1990) are less flexible than H2 because they are sublinear with respect to prey abundance. The insensitivity of the relative flexiblity of all these models to variation in predator abundances occurs because the total prey eaten they effect is determined by predator abundance in the same proportional manner as for H2. That is, just like most other two-parameter prey-dependent models, the relative flexibility of H2 and these models is similarly uninfluenced by the ratio of prey and predator abundances, in contrast to the way that all ratio- and predator-dependent models are affected (as per the contrast of H1, LR and BWL1 discussed above).

The prey-dependent Type IV model of Sokol & Howell (1981) (SH) represents an informative exception to all other two-parameter prey-dependent models in that its relative flexibility is sensitive to predator abundance. Whereas all monotonically increasing prey-dependent models only ever come up against the maximum prey abundance constraint as predator abundances increase, increasing predator abundances additionally alleviate the constraint that SH experiences uniquely due to the eventual decline of its feeding rate at high prey abundance; high predator abundances permit the total number of prey eaten to stay above the minimum-of-one-prey constraint for greater maximum prey abundances than is possible for low predator abundances given the parameter values.

The dependence of model flexibility on predator abundance emerges among the prey-dependent three-parameter models for similar reasons. For example, although the feeding rates of neither the HLB model of Hassell *et al.* (1977) nor the A2 model of Abrams (1990) decline with respect to prey abundance, increasing their *c* parameter does make their denominators more sensitive to maximum prey abundances where the minimum of one-prey-eaten-per-predator constraint comes in to play. Therefore, just as for SH, increasing predator abundances increase the number of prey eaten to allow for larger values of *c* to satisfy the minimum-of-one-prey constraint. That is, although increasing predator abundance would limit the range of *c* due to the minimum-of-one-prey constraint if all else were to be held constant, all else is not constant. Rather, high predator abundance enables a greater range of *a* values for a given value of *c* before the maximum-prey-eaten constraint is violated. This is also the reason why all predator-dependent models exhibit increasing relative flexibility as predator abundance increases even as the absolute flexibility of their respective baseline models decreases.

### 4.2 Additional aspects of experimental design

Our sensitivity analyses on the role of experimental design reinforce the inferences of our main analysis. They also speak to the likely generality of our results to additional aspects of experimental design which we did not specifically address. For the two-parameter models whose relative flexibility was insensitive to the ratio of prey and predator abundances, using arithmetic rather than logarithmic designs had little or no qualitative influence because arithmetic spacings did not alter maximum prey abundances where the constraints on the number of prey eaten are incurred. By contrast, models for which changes to spacings or the prey-eaten constraints did alter their relative flexibility were either ratio- or predator-dependent models, or were prey-dependent models whose additional (third) parameter made their flexibility sensitive to predator abundance. We conclude from this that the precise spacings of prey and predator abundances are less important from a model flexibility perspective than are their maxima and combinatorial range, but that these aspects of design become more important as the parametric complexity of the considered models increases.

Nonetheless, searching for equitable experimental designs as we did is different from searching for optimal designs for model-specific parameter uncertainty, bias, or identifiability (e.g., Moffat *et al.*, 2020; Sarnelle & Wilson, 2008; Uszko *et al.*, 2020; Zhang *et al.*, 2018). A precedence of other motivations for an experiment, such as maximizing the precision of parameter estimates, may therefore lead to different and likely model-specific conclusions about which design aspects are important. Fortunately, given our results, some aspects of experimental design may be of little consequence. For example, independent of the maximum prey abundance used, the general utility of a logarithmic spacing of prey makes intuitive sense given that, for many models, most of the action that differentiates model form occurs at low prey abundances (i.e. their derivatives with respect to *N* are greatest at low values of *N*). Intuition likewise suggests that designs should preclude total prey consumption being overwhelmed by the overall effect of interference among predators and hence that predator abundances shouldn’t be high. In this regard our results indicate that just a little variation across a range of low predator abundances is often — though far from universally — best from a relative model flexibility standpoint, just as it would be expected to be best for parameter estimation.

Our analyses did not consider questions regarding the treatment-specific distribution of experimental replicates, important though these often are given logistical constraints. All of our analyses assumed uniformly-balanced designs, the effect of which future analyses could easily assess by changing the probability of each experimental treatment when computing the Expected unit Fisher Information matrix underlying 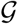 (see Box 1). We anticipate, however, that shifting replicates from lower prey and predator abundances to higher abundances will have a similar effect to that seen in the comparison of logarithmic to arithmetic spacings. Therefore, from a model flexibility standpoint alone, we expect such a shift to have a greater effect for models of high parametric complexity.

A final important aspect of experimental design that our analyses did not address was the assumed likelihood function connecting each deterministic functional-response model to an experiment’s design (i.e. the structure of the data). We assumed a Poisson likelihood and therefore that eaten prey are continually replaced, that the mean and variance of prey eaten are equal for a given combination of predator and prey abundances, and that all feeding events are independent. Model flexibility as assessed by geometric complexity may be different under alternative likelihoods such as the binomial likelihood (which would be appropriate for non-replacement designs) or the negative binomial likelihood (which allows for under- or over-dispersion). Indeed, for the binomial likelihood even the linear Holling Type I deterministic function response results in a non-linear statistical model (Novak & Stouffer, 2021), hence relative geometric complexity may be quite different for models that account for prey depletion (see *Supplementary Materials* for a comparison of Rogers’ random Type II and Type III predator models). That said, the maximum likelihood parameter estimators under Gaussian and log-Normal likelihoods are the same as under a Poisson likelihood for many — and possibly all — of the models we considered (Novak & Stouffer, 2021), so it is likely that our inferences would be little changed under these commonly assumed alternatives.

### 4.3 Model flexibility as problem and desirable property

There are many perspectives on the purpose of models and why we fit models to data. Shmueli (2010) articulates two primary axes of motivation that align well to the functional-response literature: *explanation* (where the primary motivation is to infer biologically- and statistically-significant causal associations the nature of which models are meant to characterize) and *prediction* (where the primary motivation is to best describe as yet unseen out-of-sample data).^1^ The ability to satisfy both motivations converges as the amount of data and the range of conditions the data reflect increase, thereby mirroring the inferential convergence of information criteria as sample sizes increase and cause differences in goodness-of-fit to dominate measures of model complexity. Model flexibility, and with it our analyses, would thus be irrelevant if the sample sizes of functional-response experiments were sufficiently large. Instead, sample sizes for many studies are such that model flexibility — as well as other forms of statistical and non-statistical bias (Novak & Stouffer, 2021) — preclude the conclusion that models deemed to perform best on the basis of their information-theoretic ranking are also closest to biological truth.

Empiricists fitting functional-response models to data must therefore make the explicit choice between explanation, for which criteria such as BIC and FIA are intended, and prediction, for which AIC(c), cross-validation, model-averaging, and most forms of machine learning are intended (Aho *et al.*, 2014; Höge *et al.*, 2018; Shmueli, 2010). If data is limited and explanation is the goal, then design-dependent differences in model flexibility represent a critical problem for commonly-used criteria like BIC because more flexible models will be conflated for the truth. In such contexts, it would be wise to identify the most equitable design for a specifically chosen subset of hypothesis-driven models (see also Burnham & Anderson, 2002), or, in lieu of a better reasoned solution, to use a design or multiple designs that stack the deck against leading hypotheses associated with the most flexible models. On the other hand, if data is limited and out-of-sample prediction is the goal, then model flexibility could be considered an advantage if it causes more-complex-than-true models to be selected because they are deemed to perform better, especially when the true model may not even be among those being compared (Höge *et al.*, 2018). More generally, there are clearly contexts in which ecologists wish to have generic, flexible functional-response models that merely approximate aspects of the truth in a coarse manner, be it in more descriptive statistical contexts or in theoretical contexts where the potential role of these aspects in determining qualitatively different regimes of population dynamics is of interest (e.g., AlAdwani & Saavedra, 2020; Arditi & Ginzburg, 2012; Barbier *et al.*, 2021). In these contexts, and since all models are phenomenological and hence agnostic with respect to precise mechanistic detail (as Table 1 underscores; see also Connolly *et al.*, 2017; Hart *et al.*, 2018), we consider the results of our analyses to be useful for making *a priori* choices among models given that more flexible models likely capture and exhibit a greater amount of biologically insightful variation in a more analytically tractable manner.

### 4.4 Conclusions

Several syntheses evidence that there is no single model that can characterize predator functional responses in general (Novak & Stouffer, 2021; Skalski & Gilliam, 2001; Stouffer & Novak, 2021). This is consistent with the fact that, to a large degree, the statistical models of the functional-response literature characterize aspects of predator-prey biology for which there is evidence in data, not whether specific mechanisms do or do not occur in nature (see also Connolly *et al.*, 2017). In light of the fact that functional-response data are hard to come by, our study demonstrates that a model’s functional flexibility should be considered when interpreting its performance. That said, we are not advocating for FIA as an alternative to more commonly-used information criteria; its technical nature and model-specific idiosyncrasies do not lend itself to widespread adoption or straightforward implementation (e.g., in software packages). Moreover, more fundamental issues exist that pertain to the explicit consideration of study motivation. Indeed, we submit that questions of motivation are ones that the functional-response literature as a whole needs to grapple with more directly. Even in the specific context of prediction, for example, functional-response studies rarely address explicitly what their study and their data are intending to help better predict (e.g., feeding rates or population dynamics). Valuable effort would therefore be expended in future work to consider the relationship of model flexibility to the parametric- and structural sensitivities of models when it comes to drawing inferences for population dynamics (e.g., Adamson & Morozov, 2020; Aldebert & Stouffer, 2018). Likewise, it would also be useful to clarify the relevance of model flexibility to the rapidly developing methods of scientific machine learning, including the use of symbolic regression, neural ordinary differential equations, and universal differential equations for model discovery (e.g., Bonnaffé *et al.*, 2021; Guimerà *et al.*, 2020; Martin *et al.*, 2018; Rackauckas *et al.*, 2020).

## Acknowledgments

We thank Thomas Hossie for inviting our submission and for pointing out some models we had missed, and Bryan Lynn and the two reviewers for suggestions that helped clarify the manuscript. DBS received support from the Marsden Fund Council from New Zealand Government funding, managed by Royal Society Te Apārangi (grant 16-UOC-008). MN acknowledges the support of a David and Lucile Packard Foundation grant to the Partnership for Interdisciplinary Studies of Coastal Oceans.

## Author contributions

MN led the study and wrote the first draft. Both authors contributed to the analyses and revisions.

## Code and data availability

All code has been archived at FigShare:XXXXX_◊_ and is available at https://github.com/MN:Tobeuploadeduponacceptancemarknovak/GeometricComplexity.

## Footnotes

1 A third axis, *description*, remains common in the functional-response literature and typically takes the form of fitting “non-mechanistic” polynomial models to evaluate the statistical significance of various non-linearities.

## References

Abrams, P. A. (1982). Functional responses of optimal foragers. The American Naturalist, 120, 382–390.

Abrams, P. A. (1990). The effects of adaptive-behavior on the type-2 functional-response. Ecology, 71, 877–885.

Adamson, M. W. & Morozov, A. Y. (2020). Identifying the sources of structural sensitivity in partially specified biological models. Scientific Reports, 10, 16926.

Aho, K., Derryberry, D. & Peterson, T. (2014). Model selection for ecologists: the worldviews of AIC and BIC. Ecology, 95, 631–636.

AlAdwani, M. & Saavedra, S. (2020). Ecological models: higher complexity in, higher feasibility out. Journal of The Royal Society Interface, 17, 20200607.

Aldebert, C., Nerini, D., Gauduchon, M. & Poggiale, J. C. (2016a). Does structural sensitivity alter complexity–stability relationships? Ecological Complexity, 28, 104–112.

Aldebert, C., Nerini, D., Gauduchon, M. & Poggiale, J. C. (2016b). Structural sensitivity and resilience in a predator–prey model with density-dependent mortality. Ecological Complexity, 28, 163–173.

Aldebert, C. & Stouffer, D. B. (2018). Community dynamics and sensitivity to model structure: towards a probabilistic view of process-based model predictions. Journal of The Royal Society Interface, 15, 20180741.

Andrews, J. F. (1968). A mathematical model for the continuous culture of microorganisms utilizing inhibitory substrates. Biotechnology and Bioengineering, 10, 707–723.

Arditi, R. & Akçakaya, H. R. (1990). Underestimation of mutual interference of predators. Oecologia, 83, 358–361.

Arditi, R. & Ginzburg, L. R. (1989). Coupling in predator-prey dynamics: Ratio-dependence. Journal of Theoretical Biology, 139, 311–326.

Arditi, R. & Ginzburg, L. R. (2012). How species interact: altering the standard view on trophic ecology. Oxford University Press.

Barbier, M., Wojcik, L. & Loreau, M. (2021). A macro-ecological approach to predation density-dependence. Oikos, 130, 553–570.

Beddington, J. R. (1975). Mutual interference between parasites or predators and its effect on searching efficiency. The Journal of Animal Ecology, 44, 331–340.

Bonnaffé, W., Sheldon, B. C. & Coulson, T. (2021). Neural ordinary differential equations for ecological and evolutionary time-series analysis. Methods in Ecology and Evolution, 12, 1301–1315.

Burnham, K. P. & Anderson, D. R. (2002). Model selection and multimodel inference: A practical information-theoretic approach. 2nd edn. Springer, New York.

Coelho, M. T. P., Diniz-Filho, J. & Rangel, T. F. (2019). A parsimonious view of the parsimony principle in ecology and evolution. Ecography, 42, 968–976.

Connolly, S. R., Keith, S. A., Colwell, R. K. & Rahbek, C. (2017). Process, mechanism, and modeling in macroecology. Trends in Ecology & Evolution, 32, 835–844.

Cosner, C., DeAngelis, D. L., Ault, J. S. & Olson, D. B. (1999). Effects of spatial grouping on the functional response of predators. Theoretical Population Biology, 56, 65–75.

Crowley, P. H. & Martin, E. K. (1989). Functional responses and interference within and between year classes of a dragonfly population. Journal of the North American Benthological Society, 8, 211–221

DeAngelis, D. L., Goldstein, R. A. & O’Neill, R. V. (1975). A model for trophic interaction. Ecology, 56, 881–892.

DeLong, J. P. & Uiterwaal, S. F. (2018). The FoRAGE (Functional Responses from Around the Globe in all Ecosystems) database: a compilation of functional responses for consumers and parasitoids. Knowledge Network for Biocomplexity, doi:10.5063/F17H1GTQ.

Ellison, A. M. (2004). Bayesian inference in ecology. Ecology Letters, 7, 509–520.

Evans, M. R., Grimm, V., Johst, K., Knuuttila, T., de Langhe, R., Lessells, C. M., Merz, M., O’Malley, M. A., Orzack, S. H., Weisberg, M., Wilkinson, D. J., Wolkenhauer, O. & Benton, T. G. (2013). Do simple models lead to generality in ecology? Trends in Ecology & Evolution, 28, 578–583.

Fujii, K., Holling, C. & Mace, P. (1986). A simple generalized model of attack by predators and parasites. Ecological Research, 1, 141–156.

Fussmann, G. F. & Blasius, B. (2005). Community response to enrichment is highly sensitive to model structure. Biology Letters, 1, 9–12.

Gause, G. F. (1934). The Struggle for Existence. Williams and Wilkins, Baltimore.

Grünwald, P. (2000). Model selection based on minimum description length. Journal of Mathematical Psychology, 44, 133–152.

Guimerà, R., Reichardt, I., Aguilar-Mogas, A., Massucci, F. A., Miranda, M., Pallarès, J. & Sales-Pardo, M. (2020). A bayesian machine scientist to aid in the solution of challenging scientific problems. Science Advances, 6, eaav6971.

Gutierrez, A. & Baumgärtner, J. (1984). Multitrophic models of predator-prey energetics I: Age-specific energetics models – Pea aphid *Acrythosiphon pisum* (Homoptera: Aphidae) as an example. The Canadian Entomologist, 116, 923–932.

Hart, S. P., Freckleton, R. P. & Levine, J. M. (2018). How to quantify competitive ability. Journal of Ecology, 106, 1902–1909.

Hassell, M. P., Lawton, J. H. & Beddington, J. R. (1977). Sigmoid functional responses by invertebrate predators and parasitoids. The Journal of Animal Ecology, 46, 249–262.

Hassell, M. P. & Varley, G. C. (1969). New inductive population model for insect parasites and its bearing on biological control. Nature, 223, 1133–1137.

Höge, M., Wöhling, T. & Nowak, W. (2018). A primer for model selection: The decisive role of model complexity. Water Resources Research, 54, 1688–1715.

Holling, C. S. (1959). Some characteristics of simple types of predation and parasitism. The Canadian Entomologist, 91, 385–398.

Holling, C. S. (1965). The functional response of predators to prey density and its role in mimicry and population regulation. Memoirs of the Entomological Society of Canada, 45, 3–60.

Ivlev, V. S. (1955). Experimental ecology of the feeding of fishes. Yale University Press.

Jassby, A. D. & Platt, T. (1976). Mathematical formulation of the relationship between photo-synthesis and light for phytoplankton. Limnology and Oceanography, 21, 540–547.

Jeschke, J. M., Kopp, M. & Tollrian, R. (2002). Predator functional responses: Discriminating between handling and digesting prey. Ecological Monographs, 72, 95–112.

Johnson, J. B. & Omland, K. S. (2004). Model selection in ecology and evolution. Trends in Ecology & Evolution, 19, 101–108.

Kass, R. E. & Raftery, A. E. (1995). Bayes factors. Journal of the American Statistical Association, 90, 773–795.

Kratina, P., Vos, M., Bateman, A. & Anholt, B. R. (2009). Functional responses modified by predator density. Oecologia, 159, 425–433.

Lotka, A. J. (1925). Elements of physical biology. Williams & Wilkins.

Ly, A., Marsman, M., Verhagen, J., Grasman, R. P. & Wagenmakers, E.-J. (2017). A tutorial on Fisher information. Journal of Mathematical Psychology, 80, 40 – 55.

Martin, B., Munch, S. & Hein, A. (2018). Reverse-engineering ecological theory from data. Proceedings of the Royal Society B: Biological Sciences, 285, 20180422.

Michaelis, L. & Menten, M. L. (1913). Die Kinetik der Invertinwirkung. Biochem. Z., 49, 333–369.

Moffat, H., Hainy, M., Papanikolaou, N. E. & Drovandi, C. (2020). Sequential experimental design for predator–prey functional response experiments. Journal of The Royal Society Interface, 17, 20200156.

Myung, J. I., Navarro, D. J. & Pitt, M. A. (2006). Model selection by normalized maximum likelihood. Journal of Mathematical Psychology, 50, 167–179.

Novak, M. & Stouffer, D. B. (2021). Systematic bias in studies of consumer functional responses. Ecology Letters, 24, 580–593.

Okuyama, T. (2013). On selection of functional response models: Holling’s models and more. BioControl, 58, 293–298.

Okuyama, T. & Ruyle, R. L. (2011). Solutions for functional response experiments. Acta Oecologica, 37, 512–516.

Pimm, S. L. (1982). Food webs. Springer Netherlands, Dordrecht. ISBN 978-94-009-5925-5, pp. 1–11. URL https://doi.org/10.1007/978-94-009-5925-5_1.

Pitt, M. A., Myung, I. J. & Zhang, S. (2002). Toward a method of selecting among computational models of cognition. Psychological review, 109, 472

Rackauckas, C., Ma, Y., Martensen, J., Warner, C., Zubov, K., Supekar, R., Skinner, D., Ramadhan, A. & Edelman, A. (2020). Universal differential equations for scientific machine learning. arXiv preprint arXiv:2001.04385.

Real, L. A. (1977). The kinetics of functional response. The American Naturalist, 111, 289–300.

Rissanen, J. (1978). Modeling by shortest data description. Automatica, 14, 465–471.

Rissanen, J. J. (1996). Fisher information and stochastic complexity. IEEE Transactions on Information Theory, 42, 40–47.

Rogers, D. (1972). Random search and insect population models. Journal of Animal Ecology, 41, 369–383.

Rosenzweig, M. L. (1971). Paradox of enrichment: Destabilization of exploitation ecosystems in ecological time. Science, 171, 385–387.

Ruxton, G., Gurney, W. & De Roos, A. (1992). Interference and generation cycles. Theoretical Population Biology, 42, 235–253.

Sarnelle, O. & Wilson, A. E. (2008). Type III functional response in Daphnia. Ecology, 89, 1723–1732

Schenk, D., Bersier, L.-F. & Bacher, S. (2005). An experimental test of the nature of predation: neither prey- nor ratio-dependent. Journal of Animal Ecology, 74, 86–91.

Shmueli, G. (2010). To explain or to predict? Statististical Science, 25, 289–310.

Skalski, G. T. & Gilliam, J. F. (2001). Functional responses with predator interference: Viable alternatives to the Holling Type II model. Ecology, 82, 3083–3092.

Sokol, W. & Howell, J. A. (1981). Kinetics of phenol oxidation by washed cells. Biotechnology and Bioengineering, 23, 2039–2049.

Stouffer, D. B. & Novak, M. (2021). Hidden layers of density dependence in consumer feeding rates. Ecology Letters, 24, 520–532.

Sutherland, W. J. (1983). Aggregation and the ‘ideal free’ distribution. Journal of Animal Ecology, 52, 821–828.

Tostowaryk, W. (1972). The effect of prey defense on the functional response of *Podisus modestus* (Hemiptera: Pentatomidae) to densities of the sawflies *Neodiprion swainei* and *N. pratti banksianae* (Hymenoptera: Neodiprionidae). The Canadian Entomologist, 104, 61–69.

Tyutyunov, Y., Titova, L. & Arditi, R. (2008). Predator interference emerging from trophotaxis in predator–prey systems: An individual-based approach. Ecological Complexity, 5, 48–58.

Uszko, W., Diehl, S. & Wickman, J. (2020). Fitting functional response surfaces to data: a best practice guide. Ecosphere, 11, e03051.

Volterra, V. (1926). Fluctuations in the abundance of a species considered mathematically. Nature, 118, 558–560.

Watt, K. (1959). A mathematical model for the effect of densities of attacked and attacking species on the number attacked. The Canadian Entomologist, 91, 129–144.

Whitehead, A. N. (1919). The concept of nature. Convergence, 62, 79.

Wolfram Research Inc. (2020). Mathematica, v.12.1. Champaign, IL. URL https://www.wolfram.com/mathematica.

Zhang, J. F., E., P. N., Theodore, K. & C., D. C. (2018). Optimal experimental design for predator–prey functional response experiments. Journal of The Royal Society Interface, 15, 20180186.

