## Supplementary Materials for "Geometric complexity and the information-theoretic comparison of functional-response models"

### Contents

**S1 Original model parameterizations** **2**

**S2 Fisher Information matrix** **3**

**S3 Model identifiability** **4**

**S4 Experimental designs** **7**

**S5 Results for fixed logarithmic designs** **8**

**S6 Results for arithmetic experimental designs** **8**

**S7 Results for logarithmic designs with $\mathbb{E}[F(N_{max}, P, \theta)PT] \geq 1/10$ constraint** **17**

**S8 Results for logarithmic designs with $\mathbb{E}[F(N, P, \theta)PT] \leq 10N_{max}$ constraint** **21**

**S9 Geometric complexity in non-replacement experiments** **25**

**References** **28**

### S1 Original model parameterizations

Table S1: Original formulations for the models which we re-parameterized (with the exception of the Hyperbolic tangent, which is written here in an alternative formulation). Note that we divide the models of Barbier *et al.* (2021) by $P$ because their formulations describe population-level feeding rates whereas all others describe per-predator feeding rates.

| Name | Original | Reference |
|---|---|---|
| **One-parameter models ($k = 1$)** | | |
| Barbier–Wojcik–Loreau I | $a\sqrt{NP}/P$ | Barbier *et al.* (2021) |
| **Two-parameter models ($k = 2$)** | | |
| Abrams I | $\sqrt{\frac{qcN}{d(1+chN)}}$ | Abrams (1990) |
| Abrams III | $\frac{\sqrt{N}}{2\sqrt{uH}+h\sqrt{N}}$ | Abrams (1990) |
| Hyperbolic tangent | $\frac{1}{b}\frac{\exp[2abN]+1}{\exp[2abN]-1}$ | Jassby & Platt (1976) |
| Sokol–Howell | $\frac{aN}{b+N^2}$ | Sokol & Howell (1981) |
| **Three-parameter models ($k = 3$)** | | |
| Abrams II | $\frac{cN}{1+h_{min}cN+\sqrt{\frac{cN}{a}(1+ch_{min}N)}}$ | Abrams (1990) |
| Hassell–Lawton–Beddington | $\frac{aN^2}{1+cN+abN^2}$ | Hassell *et al.* (1977) |
| Monod–Haldane | $\frac{aN}{b+N+N^2/c}$ | Andrews (1968) |
| Tostowaryk | $\frac{aN}{1+abN+acN^3}$ | Tostowaryk (1972) |
| Fujii–Holling–Mace | $\frac{1}{b+1/(aN\exp[dN])}$ | Fujii *et al.* (1986) |
| Steady State Satiation | $\frac{1+a(b+c)N-\sqrt{1+aN(2(b+c)+aN(b-c)^2)}}{2abcN}$ | Jeschke *et al.* (2002) |
| Tyutyunov–Titova–Arditi | $\frac{aN}{P/P_0+\exp[-P/P_0]+ahN}$ | Tyutyunov *et al.* (2008) |
| Barbier–Wojcik–Loreau II | $aN^uP^v/P$ | Barbier *et al.* (2021) |

### S2 Fisher Information matrix

As described in Box 1, the unit Fisher Information matrix $I_M(\theta)$ used to compute geometric complexity is a $k \times k$ matrix comprising the expected values of the second-order derivatives of the model's negative log-likelihood function with respect to each of its $k$ parameters (Pitt *et al.*, 2002; Rissanen, 1996). As an example, for a one-parameter model with $n = 2$ experimental treatments this "matrix" is given by

$$\mathcal{I}_M(\theta) = -\frac{1}{2}\,\mathbb{E}\left[\frac{\partial}{\partial\theta}\frac{\partial}{\partial\theta}\ln\mathcal{L}(\theta|\,\{y_1, y_2\})\right] \tag{S1}$$

which for discrete outcomes is equivalent to

$$\mathcal{I}_M(\theta) = -\frac{1}{2}\sum_{\{y_1,y_2\}} p\left(\{y_1, y_2\}\,|\theta\right)\left[\frac{\partial}{\partial\theta}\frac{\partial}{\partial\theta}\ln\mathcal{L}(\theta|\,\{y_1, y_2\})\right], \tag{S2}$$

and the sum is across the Fisher information for all potentially observable outcomes $\{y_1, y_2\}$ weighted by their probability of being observed given the value of $\theta$. When assuming a Poisson statistical model for the observations, we can rewrite Eq. (S1) as

$$\mathcal{I}_{\mathcal{M}}(\theta) = -\frac{1}{2}\,\mathbb{E}\left[\frac{\partial}{\partial\theta}\frac{\partial}{\partial\theta}\left(-\prod_{i=1}^{2}\ln\left(y_i!\right) + \sum_{i=1}^{2}\left(\ln(\lambda_i)y_i - \lambda_i\right)\right)\right], \tag{S3}$$

recalling that $\lambda_i$ is a function of parameter $\theta$ as determined by the functional-response model $M$. The first term from the log-likelihood drops out upon differentiating because it does not depend on $\theta$, leaving

$$\mathcal{I}_{\mathcal{M}}(\theta) = -\frac{1}{2}\,\mathbb{E}\left[\frac{\partial}{\partial\theta}\frac{\partial}{\partial\theta}\left(\sum_{i=1}^{2}\left(\ln(\lambda_i)y_i - \lambda_i\right)\right)\right], \tag{S4}$$

which because of linearity of the expected value and differentiation is equivalent to

$$\mathcal{I}_{\mathcal{M}}(\theta) = -\frac{1}{2}\sum_{i=1}^{2}\mathbb{E}\left[\frac{\partial}{\partial\theta}\frac{\partial}{\partial\theta}\left(\ln(\lambda_i)y_i - \lambda_i\right)\right]. \tag{S5}$$

This implies that the Fisher information of multiple independent outcomes is the average of the Fisher information computed separately for each random outcome. Though this example focuses on a hypothetical model with a single parameter (i.e. $k = 1$), the same holds true for more complex models where $\theta$ is a vector because the expectation is calculated separately for each element in the Fisher information matrix (Pitt *et al.*, 2002; Rissanen, 1996).

### S3 Model identifiability

When fitting any model to data, it is vitally important that the experimental design allow inference of all underlying model parameters (Beck & Arnold, 1977; Cole, 2020; Shapiro, 1986). For example, one cannot fit a linear regression model of the form $y = \alpha + \beta x$ without variation in the predictor variable $x$. Likewise, one cannot uniquely identify the parameters of a linear regression model of the form $y = \alpha_1 + \alpha_2 + \beta x$, because any equivalent sum of $\alpha_1 + \alpha_2$ will produce equivalent results.

For our purposes, geometric flexibility $\mathcal{G}$ vanishes when any two parameters capture redundant information and are not uniquely identifiable statistically. In order to evaluate the identifiability of the parameters within the various functional-response models for our synthetic experimental designs, we followed the following process. For simplicity, we will use the three-parameter Beddington–DeAngelis (BD) model (Beddington, 1975; DeAngelis *et al.*, 1975) for our example.

First, we computed the partial derivatives of the predicted number of eaten prey given the functional-response model (i.e. $\lambda = F(N, P, \theta)PT$) with respect to the model parameters. For the BD model, this produces

$$\frac{\partial \lambda}{\partial a} = \frac{NPT(1 + c(P-1))}{(1 + abN + c(P-1))^2} \tag{S6}$$

$$\frac{\partial \lambda}{\partial b} = -\frac{a^2N^2PT}{(1 + abN + c(P-1))^2} \tag{S7}$$

$$\frac{\partial \lambda}{\partial c} = -\frac{aNP(P-1)T}{(1 + abN + c(P-1))^2} \,. \tag{S8}$$

We then construct a matrix $\mathcal{Z}$ arranged with the model parameters along the columns, a row for each of the $N \times P$ treatment combinations in the experimental design, and whose elements are the values of these partial derivatives with $N$ and $P$ replaced by their corresponding values. For example, given an experimental design that features two prey levels, $N \in \{1, 2\}$, and two predator levels, $P \in \{1, 2\}$, this matrix is

$$\mathcal{Z} = \begin{bmatrix} \frac{T}{(1+ab)^2} & -\frac{a^2T}{(1+ab)^2} & 0 \\ \frac{2(1+c)T}{(1+ab+c)^2} & -\frac{2a^2T}{(1+ab+c)^2} & -\frac{2aT}{(1+ab+c)^2} \\ \frac{2T}{(1+2ab)^2} & -\frac{4a^2T}{(1+2ab)^2} & 0 \\ \frac{4(1+c)T}{(1+2ab+c)^2} & -\frac{8a^2T}{(1+2ab+c)^2} & -\frac{4aT}{(1+2ab+c)^2} \end{bmatrix} . \tag{S9}$$

In order for all parameters to be identifiable, the rank of $\mathcal{Z}$ must be equal to the number of free parameters (Beck & Arnold, 1977; Cole, 2020; Shapiro, 1986). In the case of Eq. (S9), the rank of $\mathcal{Z}$ is 3 implying that the experimental design is sufficient to inform estimates of $a$, $b$, and $c$ in the BD model.

When the sets of partial derivatives in any two (or more) columns are linearly dependent, this implies that the corresponding parameters are not identifiable. For example, the matrix that arises with an experimental design that features two prey levels, $N \in \{1, 2\}$, and only one predator level, $P \in \{1\}$, is

$$\mathcal{Z} = \begin{bmatrix} \frac{T}{(1+ab)^2} & -\frac{a^2T}{(1+ab)^2} & 0 \\ \frac{2T}{(1+2ab)^2} & -\frac{4a^2T}{(1+2ab)^2} & 0 \end{bmatrix}, \tag{S10}$$

which implies that the parameter $c$ is non-identifiable because the third column in Eq. (S10) is linearly dependent on *both* the first and second columns. In this example, given the experimental design, the non-identifiable nature of parameter $c$ should not be all that surprising: $c$ is meant to capture interference between predators, but with only ever a single predator individual (i.e. $P = 1$) there can be no interference effect to estimate.

That said, the linear dependence and non-identifiability of $c$ is not simply a product of the chosen predator abundance level. Indeed, parameter $c$ will never be identifiable in an experimental design that features only one predator level. With an experimental design that features two prey levels, $N \in \{1, 2\}$, and only one predator level, $P \in \{2\}$, the matrix is

$$\mathcal{Z} = \begin{bmatrix} \frac{2(1+c)T}{(1+ab+c)^2} & -\frac{2a^2T}{(1+ab)^2} & -\frac{2aT}{(1+ab+c)^2} \\ \frac{4(1+c)T}{(1+2ab+c)^2} & -\frac{8a^2T}{(1+2ab)^2} & -\frac{4aT}{(1+2ab+c)^2} \end{bmatrix}, \tag{S11}$$

and the third column is again linearly dependent on the first by a factor of $-\frac{1+c}{a}$. This is because parameter identifiability also requires sufficient *potential* variation along the direction determined by each parameter, and without appropriate treatment levels there is no "slope" that can be robustly inferred (Beck & Arnold, 1977; Cole, 2020; Shapiro, 1986).

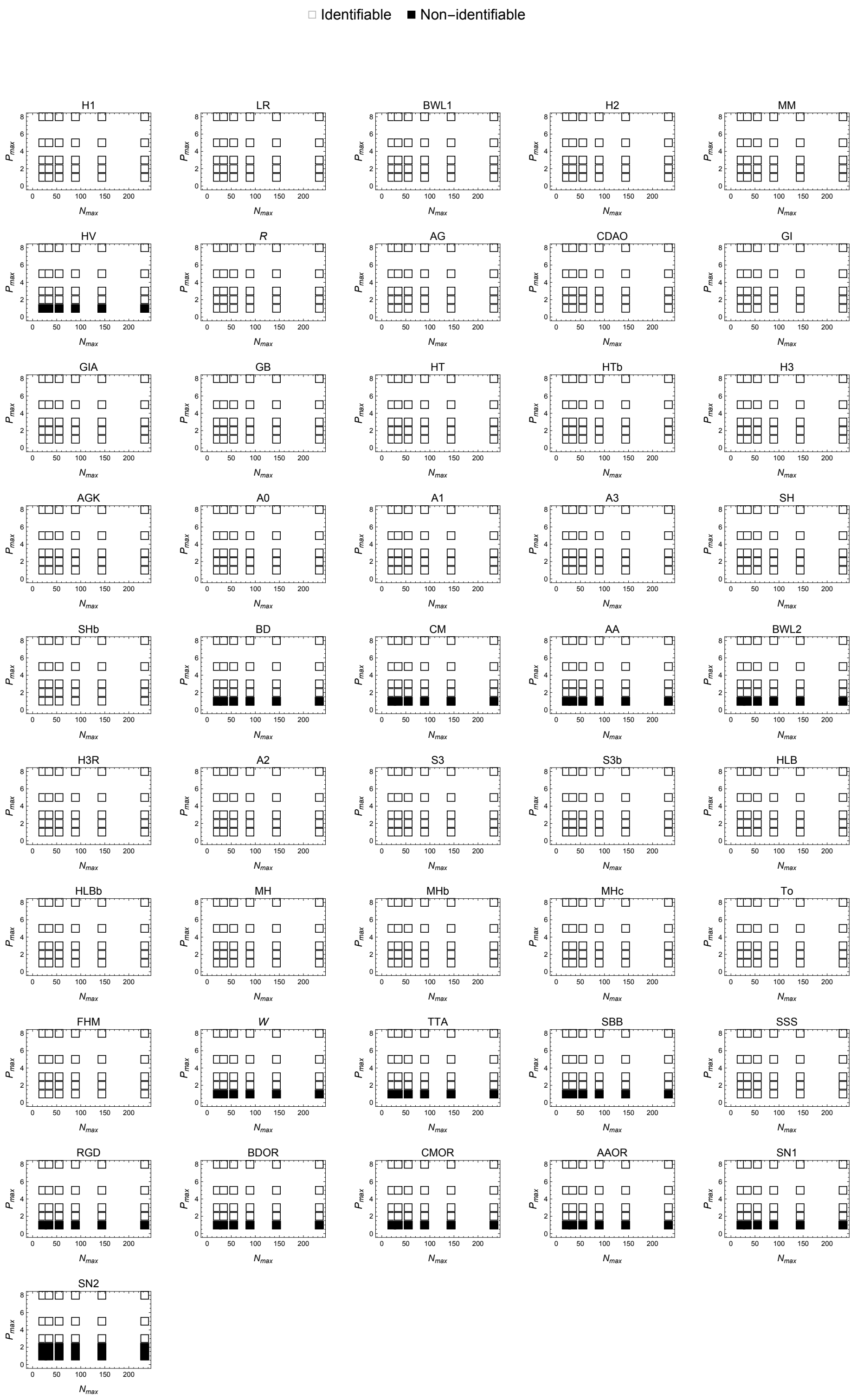

Figure S1: Model identifiability as a function of $N_{max}$ and $P_{max}$.

### S4 Experimental designs

Each nested pair of vectors reflects the abundances of the prey (first vector) and of the predator (second vector) for a given experimental design. Variable designs vary the maximum number of prey and predator individuals ($N_{max}$ and $P_{max}$) by correspondingly varying the number of prey and predator levels ($L_N$ and $L_P$). Fixed designs vary only the number of prey and predator levels ($L_N$ and $L_P$) by keeping the maximum number of prey and predator individuals constant. The arithmetic designs consist of uniform (rather than logarithmic) spacings and are variable designs.

#### S4.1 Logarithmic designs

##### S4.1.1 Variable designs

{{3, 4, 7, 13, 21}, {1}}
{{3, 4, 7, 12, 20, 34}, {1}}
{{3, 4, 7, 12, 20, 33, 55}, {1}}
{{3, 4, 7, 12, 20, 32, 54, 89}, {1}}
{{3, 4, 7, 12, 19, 32, 53, 87, 144}, {1}}
{{3, 4, 7, 12, 19, 32, 52, 86, 141, 233}, {1}}

{{3, 4, 7, 13, 21}, {1, 2}}
{{3, 4, 7, 12, 20, 34}, {1, 2}}
{{3, 4, 7, 12, 20, 33, 55}, {1, 2}}
{{3, 4, 7, 12, 20, 32, 54, 89}, {1, 2}}
{{3, 4, 7, 12, 19, 32, 53, 87, 144}, {1, 2}}
{{3, 4, 7, 12, 19, 32, 52, 86, 141, 233}, {1, 2}}

{{3, 4, 7, 13, 21}, {1, 2, 3}}
{{3, 4, 7, 12, 20, 34}, {1, 2, 3}}
{{3, 4, 7, 12, 20, 33, 55}, {1, 2, 3}}
{{3, 4, 7, 12, 20, 32, 54, 89}, {1, 2, 3}}
{{3, 4, 7, 12, 19, 32, 53, 87, 144}, {1, 2, 3}}
{{3, 4, 7, 12, 19, 32, 52, 86, 141, 233}, {1, 2, 3}}

{{3, 4, 7, 13, 21}, {1, 2, 3, 5}}
{{3, 4, 7, 12, 20, 34}, {1, 2, 3, 5}}
{{3, 4, 7, 12, 20, 33, 55}, {1, 2, 3, 5}}
{{3, 4, 7, 12, 20, 32, 54, 89}, {1, 2, 3, 5}}
{{3, 4, 7, 12, 19, 32, 53, 87, 144}, {1, 2, 3, 5}}
{{3, 4, 7, 12, 19, 32, 52, 86, 141, 233}, {1, 2, 3, 5}}

{{3, 4, 7, 13, 21}, {1, 2, 3, 5, 8}}
{{3, 4, 7, 12, 20, 34}, {1, 2, 3, 5, 8}}
{{3, 4, 7, 12, 20, 33, 55}, {1, 2, 3, 5, 8}}
{{3, 4, 7, 12, 20, 32, 54, 89}, {1, 2, 3, 5, 8}}
{{3, 4, 7, 12, 19, 32, 53, 87, 144}, {1, 2, 3, 5, 8}}
{{3, 4, 7, 12, 19, 32, 52, 86, 141, 233}, {1, 2, 3, 5, 8}}

##### S4.1.2 Fixed designs

{{3, 8, 25, 76, 233}, {1}}
{{3, 6, 16, 39, 95, 233}, {1}}
{{3, 6, 12, 25, 52, 110, 233}, {1}}
{{3, 5, 9, 18, 34, 65, 123, 233}, {1}}
{{3, 5, 8, 14, 25, 43, 76, 133, 233}, {1}}
{{3, 4, 7, 12, 19, 32, 52, 86, 141, 233}, {1}}

{{3, 8, 25, 76, 233}, {1, 5}}
{{3, 6, 16, 39, 95, 233}, {1, 5}}
{{3, 6, 12, 25, 52, 110, 233}, {1, 5}}
{{3, 5, 9, 18, 34, 65, 123, 233}, {1, 5}}
{{3, 5, 8, 14, 25, 43, 76, 133, 233}, {1, 5}}
{{3, 4, 7, 12, 19, 32, 52, 86, 141, 233}, {1, 5}}

{{3, 8, 25, 76, 233}, {1, 2, 5}}
{{3, 6, 16, 39, 95, 233}, {1, 2, 5}}
{{3, 6, 12, 25, 52, 110, 233}, {1, 2, 5}}
{{3, 5, 9, 18, 34, 65, 123, 233}, {1, 2, 5}}
{{3, 5, 8, 14, 25, 43, 76, 133, 233}, {1, 2, 5}}
{{3, 4, 7, 12, 19, 32, 52, 86, 141, 233}, {1, 2, 5}}

{{3, 8, 25, 76, 233}, {1, 2, 3, 5}}
{{3, 6, 16, 39, 95, 233}, {1, 2, 3, 5}}
{{3, 6, 12, 25, 52, 110, 233}, {1, 2, 3, 5}}
{{3, 5, 9, 18, 34, 65, 123, 233}, {1, 2, 3, 5}}
{{3, 5, 8, 14, 25, 43, 76, 133, 233}, {1, 2, 3, 5}}
{{3, 4, 7, 12, 19, 32, 52, 86, 141, 233}, {1, 2, 3, 5}}

#### S4.2 Arithmetic designs

{{3, 8, 12, 16, 21}, {1}}
{{3, 9, 15, 22, 28, 34}, {1}}
{{3, 12, 20, 29, 38, 46, 55}, {1}}
{{3, 15, 28, 40, 52, 64, 77, 89}, {1}}
{{3, 21, 38, 56, 74, 91, 109, 126, 144}, {1}}
{{3, 29, 54, 80, 105, 131, 156, 182, 207, 233}, {1}}

{{3, 8, 12, 16, 21}, {1, 2}}
{{3, 9, 15, 22, 28, 34}, {1, 2}}
{{3, 12, 20, 29, 38, 46, 55}, {1, 2}}
{{3, 15, 28, 40, 52, 64, 77, 89}, {1, 2}}
{{3, 21, 38, 56, 74, 91, 109, 126, 144}, {1, 2}}
{{3, 29, 54, 80, 105, 131, 156, 182, 207, 233}, {1, 2}}

{{3, 8, 12, 16, 21}, {1, 2, 3}}
{{3, 9, 15, 22, 28, 34}, {1, 2, 3}}
{{3, 12, 20, 29, 38, 46, 55}, {1, 2, 3}}
{{3, 15, 28, 40, 52, 64, 77, 89}, {1, 2, 3}}
{{3, 21, 38, 56, 74, 91, 109, 126, 144}, {1, 2, 3}}
{{3, 29, 54, 80, 105, 131, 156, 182, 207, 233}, {1, 2, 3}}

{{3, 8, 12, 16, 21}, {1, 2, 4, 5}}
{{3, 9, 15, 22, 28, 34}, {1, 2, 4, 5}}
{{3, 12, 20, 29, 38, 46, 55}, {1, 2, 4, 5}}
{{3, 15, 28, 40, 52, 64, 77, 89}, {1, 2, 4, 5}}
{{3, 21, 38, 56, 74, 91, 109, 126, 144}, {1, 2, 4, 5}}
{{3, 29, 54, 80, 105, 131, 156, 182, 207, 233}, {1, 2, 4, 5}}

{{3, 8, 12, 16, 21}, {1, 3, 4, 6, 8}}
{{3, 9, 15, 22, 28, 34}, {1, 3, 4, 6, 8}}
{{3, 12, 20, 29, 38, 46, 55}, {1, 3, 4, 6, 8}}
{{3, 15, 28, 40, 52, 64, 77, 89}, {1, 3, 4, 6, 8}}
{{3, 21, 38, 56, 74, 91, 109, 126, 144}, {1, 3, 4, 6, 8}}
{{3, 29, 54, 80, 105, 131, 156, 182, 207, 233}, {1, 3, 4, 6, 8}}

### S5 Results for fixed logarithmic designs

Figures S2–S5 show the effect of varying only the number of prey and predator levels ($L_N$ and $L_P$) by keeping the maximum number of prey and predator individuals constant. For these *fixed* designs, we set the maximum possible values of $N_{max}$ and $P_{max}$ to be 233 prey and 5 predator individuals (based on results observed for the *variable* designs, as presented in the main text) and then varied $L_N$ between 5 and 10 prey levels and $L_P$ between 1 and 4 predator levels. As for the *variable* designs presented in the main text, the spacing of prey (and predator) abundances followed a logarithmic series using the golden ratio as the logarithmic base.

### S6 Results for arithmetic experimental designs

Figures S6–S9 present the results for arithmetic experimental designs that used a uniform (rather than logarithmic) spacing of prey and predator abundances and varied the maximum number of prey and predator individuals ($N_{max}$ and $P_{max}$) by correspondingly varying the number of prey and predator levels ($L_N$ and $L_P$) (i.e. including additional, higher abundance levels to smaller designs).

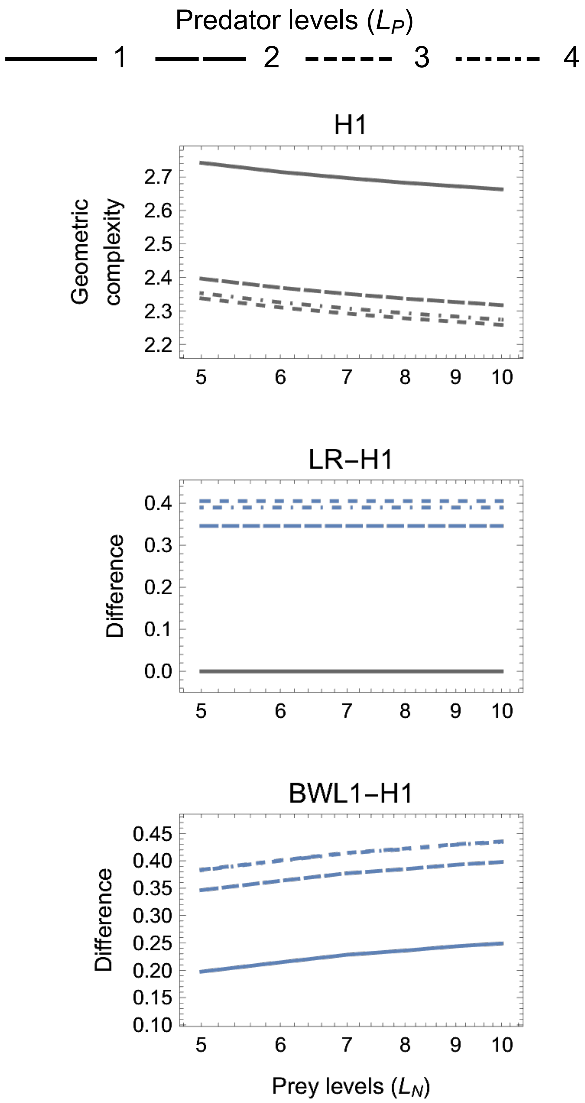

Figure S2: *First panel*: Absolute and relative geometric complexity for the one-parameter ($k = 1$) models as a function of the experiment's number of treatment levels ($L_N$ and $L_P$).

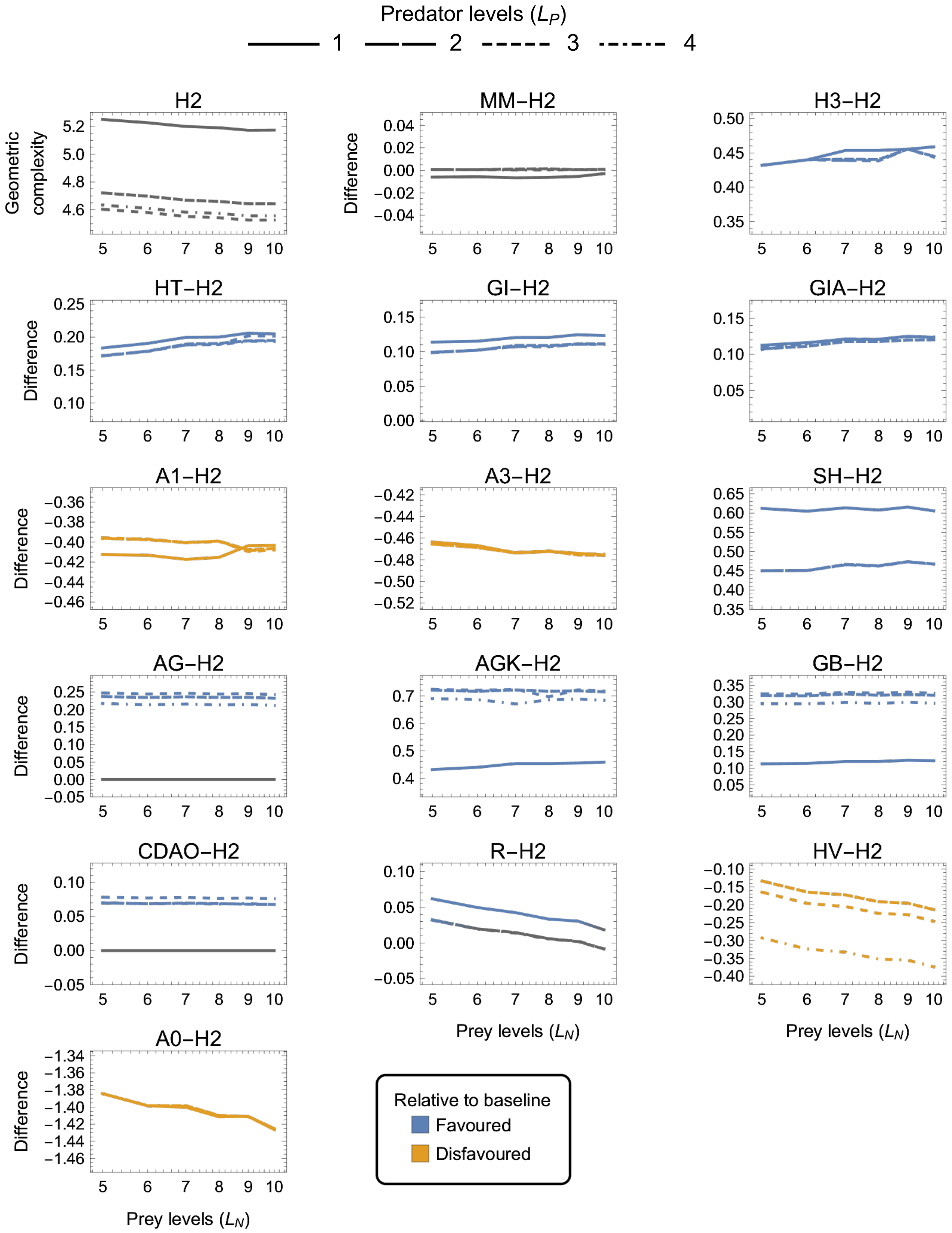

Figure S3: Absolute and relative geometric complexity for the two-parameter ($k = 2$) models as a function of the experiment's number of treatment levels ($L_N$ and $L_P$).

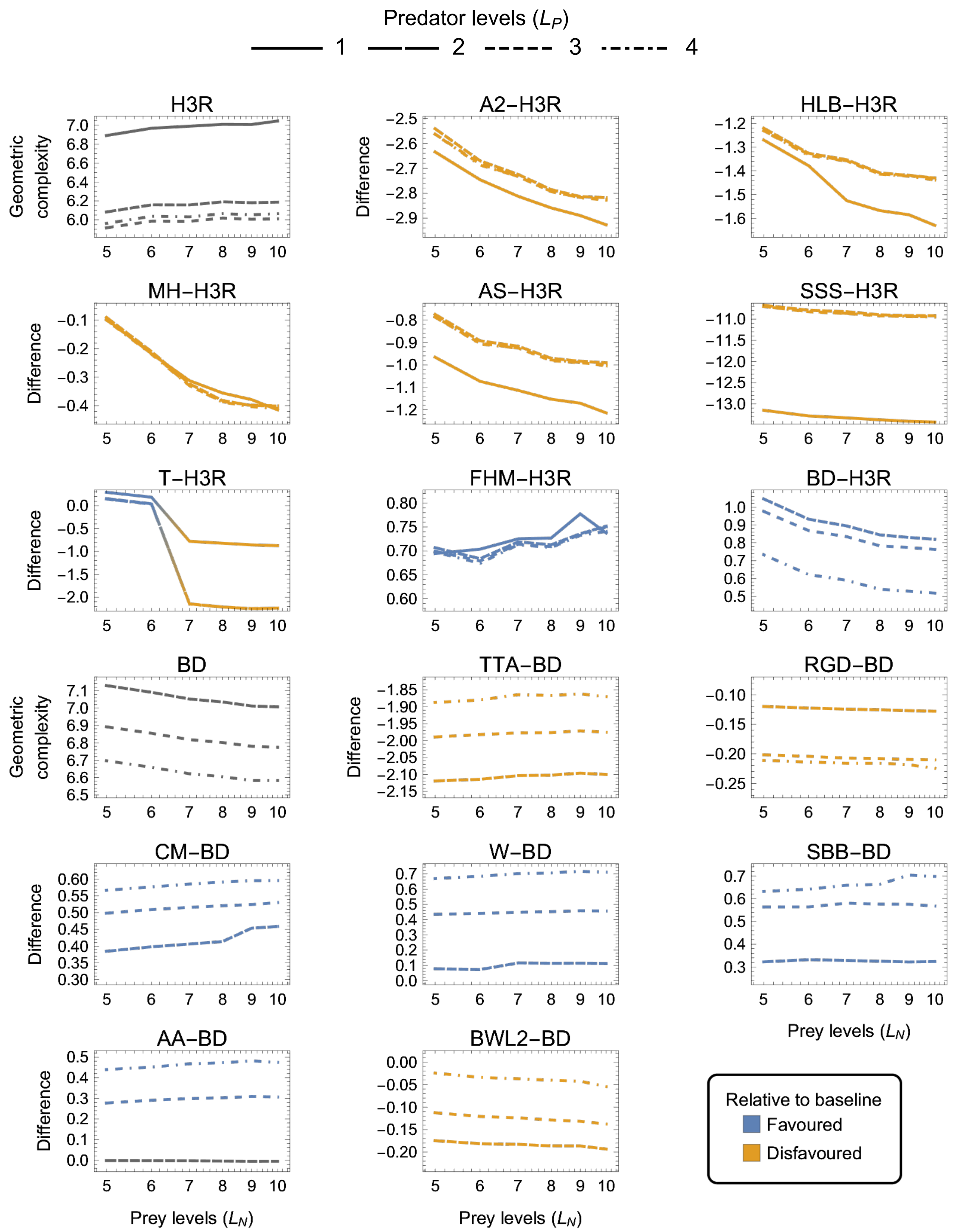

Figure S4: Absolute and relative geometric complexity for the three-parameter ($k = 3$) models as a function of the experiment's number of treatment levels ($L_N$ and $L_P$).

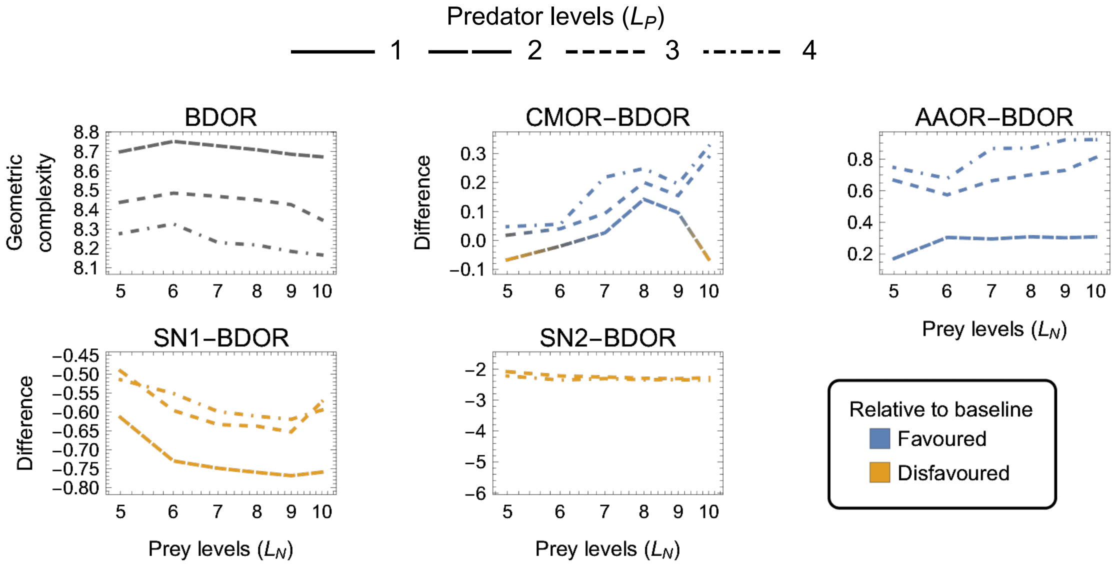

Figure S5: Absolute and relative geometric complexity for the four-parameter ($k = 4$) models as a function of the experiment's number of treatment levels ($L_N$ and $L_P$).

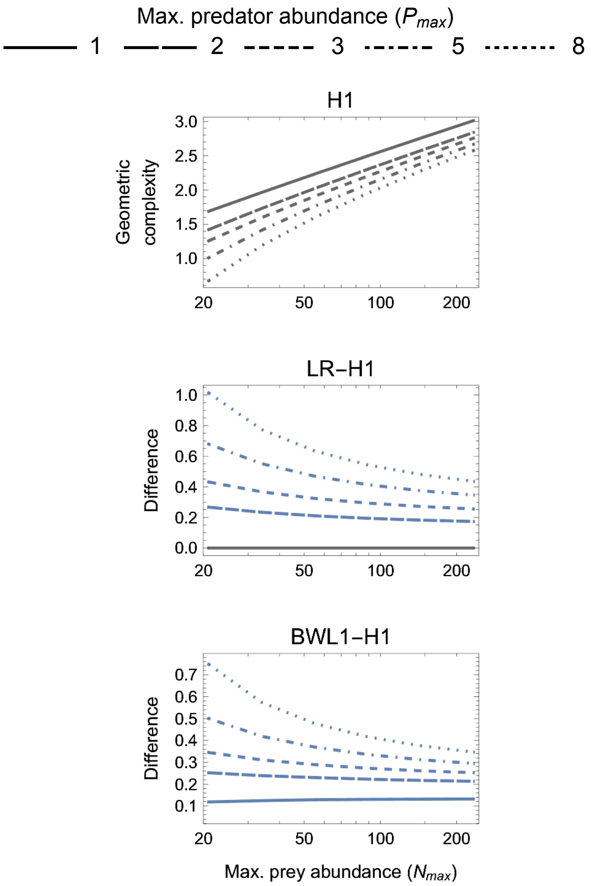

Figure S6: Absolute and relative geometric complexity for the one-parameter ($k = 1$) models with arithmetic spacing of prey and predator abundances.

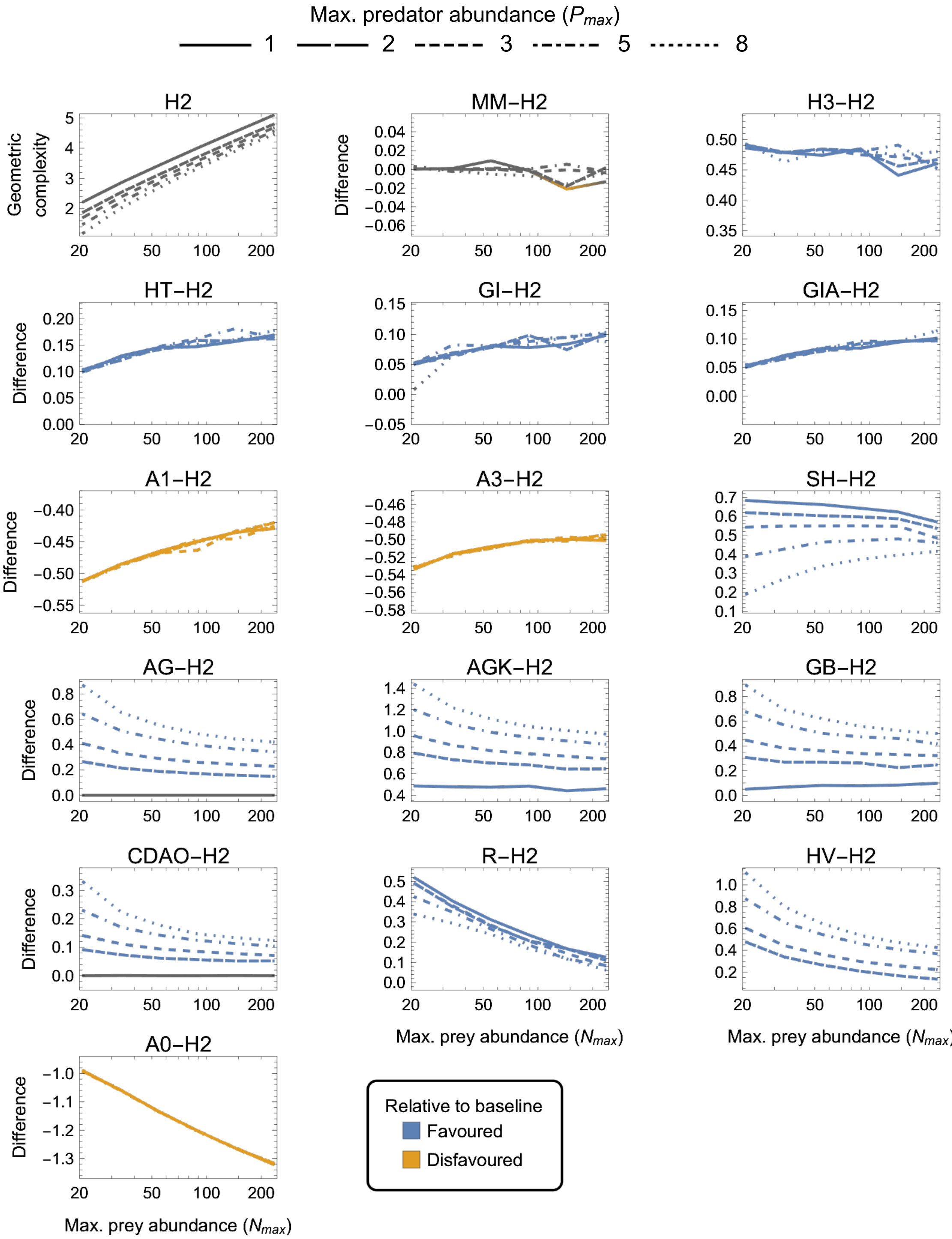

Figure S7: Absolute and relative geometric complexity for the two-parameter ($k = 2$) models with arithmetic spacing of prey and predator abundances.

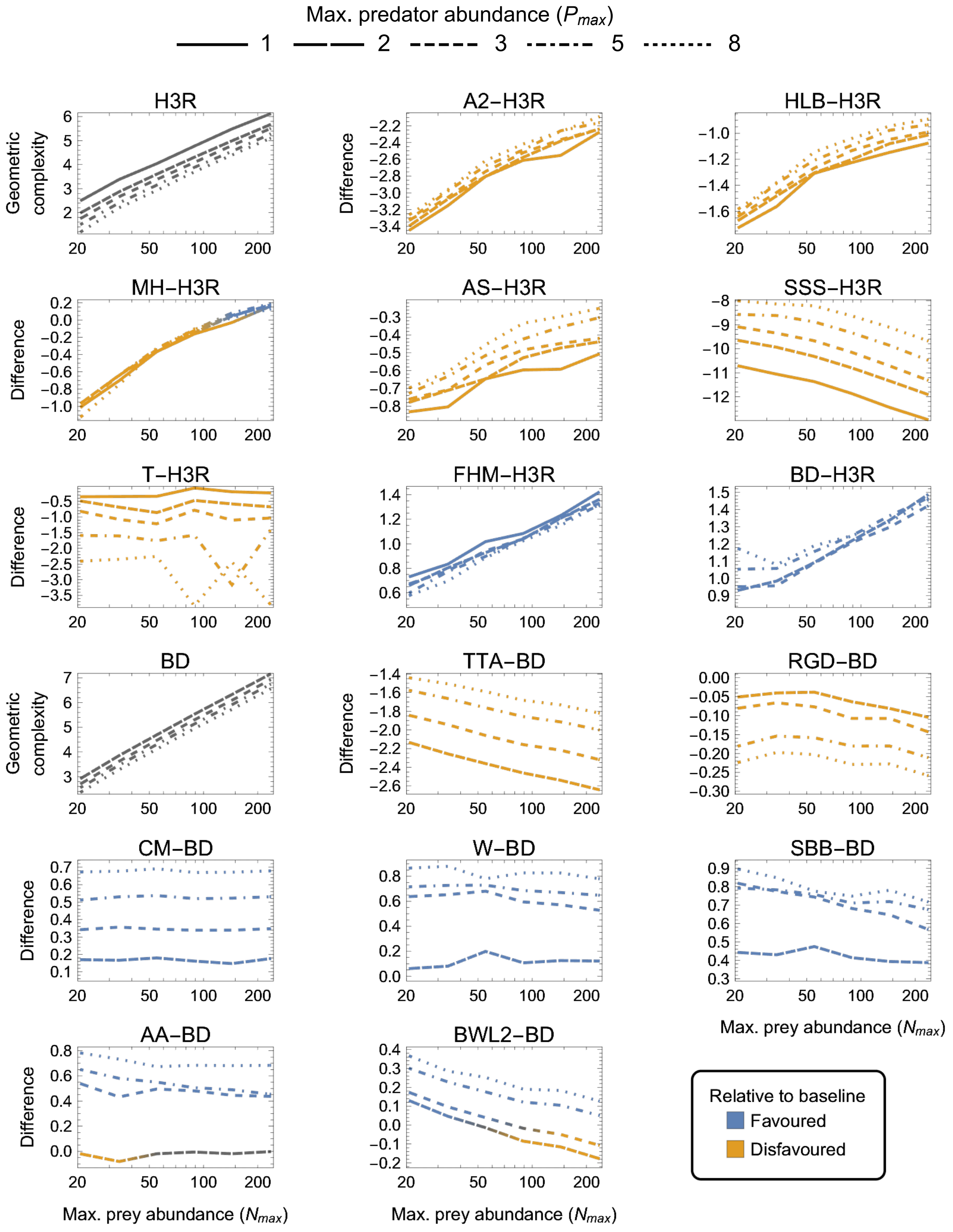

Figure S8: Absolute and relative geometric complexity for the three-parameter ($k = 3$) models with arithmetic spacing of prey and predator abundances..

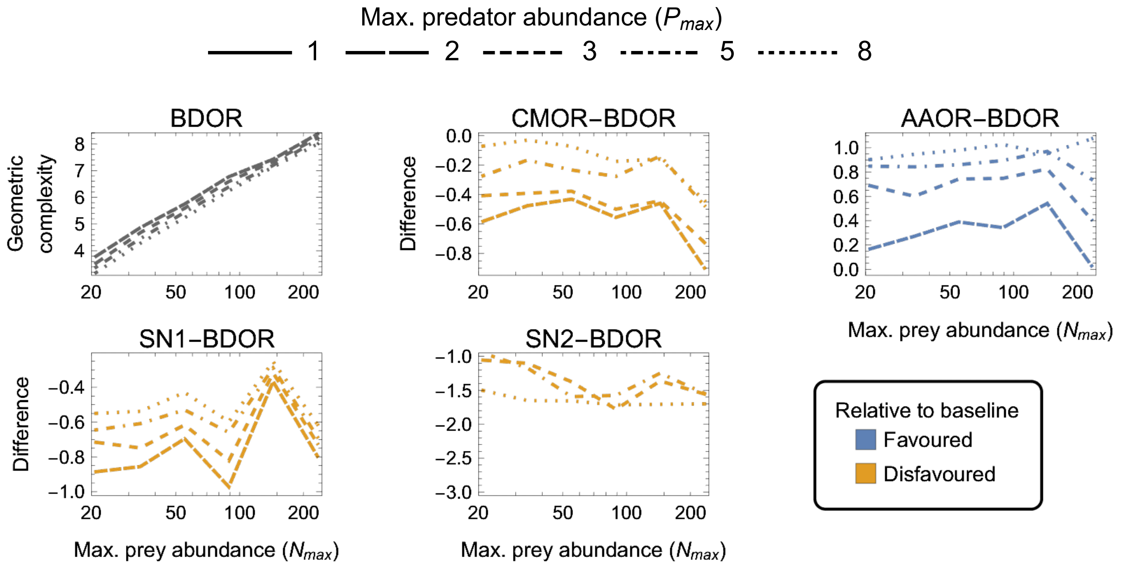

Figure S9: Absolute and relative geometric complexity for the four-parameter ($k = 4$) models with arithmetic spacing of prey and predator abundances.

### S7 Results for logarithmic designs with $\mathbb{E}[F(N_{max}, P, \theta)PT] \geq 1/10$ constraint

Figures S10–S13 present the results for logarithmic experimental designs on which the constraint on the minimum expected number of eaten prey was decreased by an order of magnitude from $\mathbb{E}[F(N_{max}, P, \theta)PT] \geq 1$ to $\mathbb{E}[F(N_{max}, P, \theta)PT] \geq 1/10$.

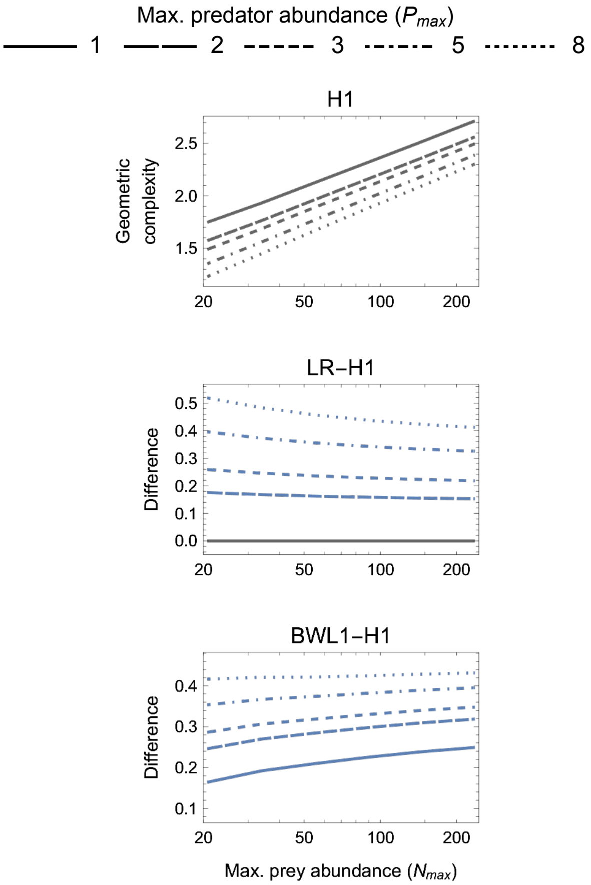

Figure S10: Absolute and relative geometric complexity for the one-parameter ($k = 1$) models with the constraint on the expected minimum number of eaten prey in the $N_{max}$ treatments being decreased by an order of magnitude (i.e. $\mathbb{E}[F(N_{max}, P, \theta)PT] \geq 1/10$).

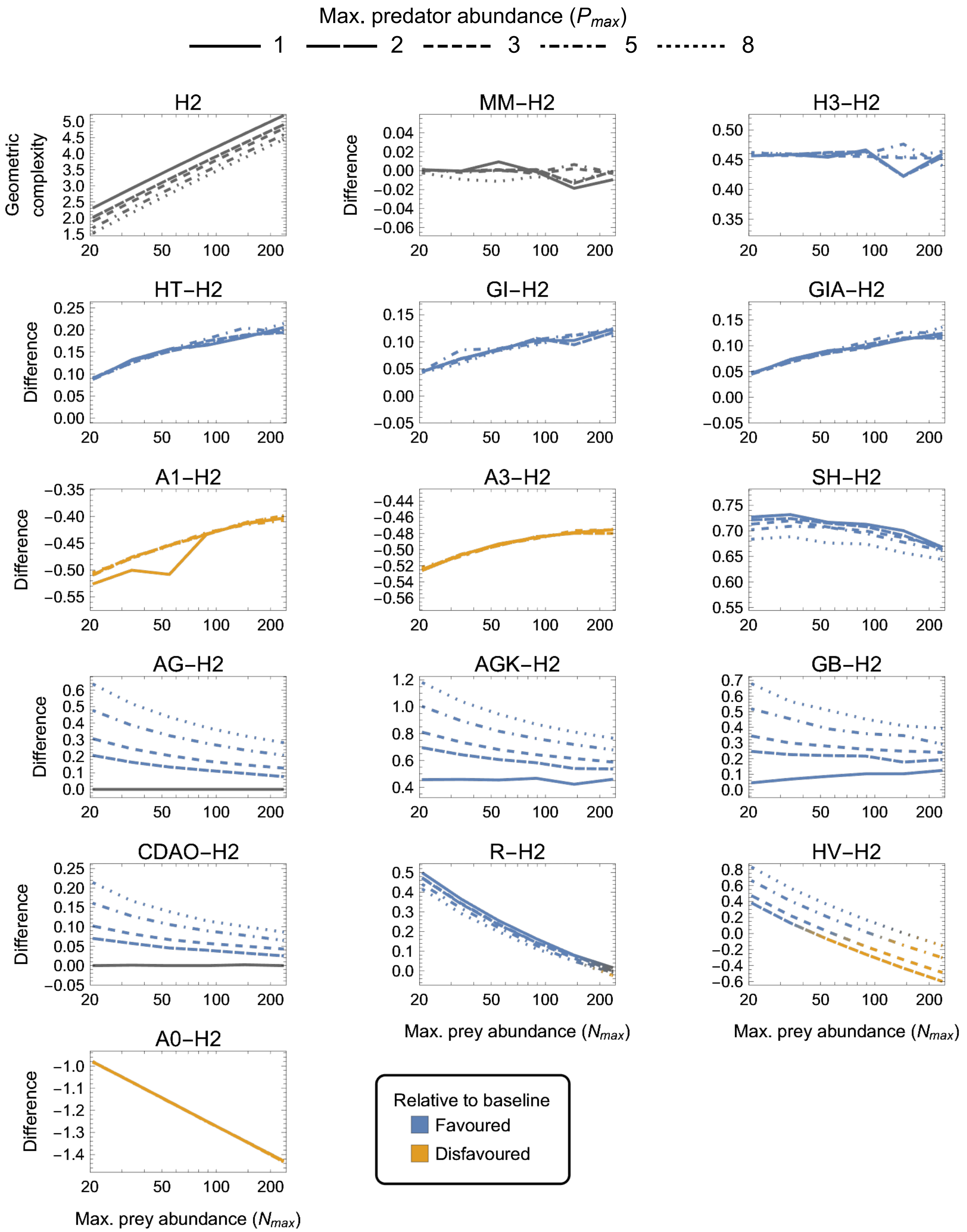

Figure S11: Absolute and relative geometric complexity for the two-parameter ($k = 2$) models with the constraint on the expected minimum number of eaten prey in the $N_{max}$ treatments being decreased by an order of magnitude (i.e. $\mathbb{E}[F(N_{max}, P, \theta)PT] \geq 1/10$).

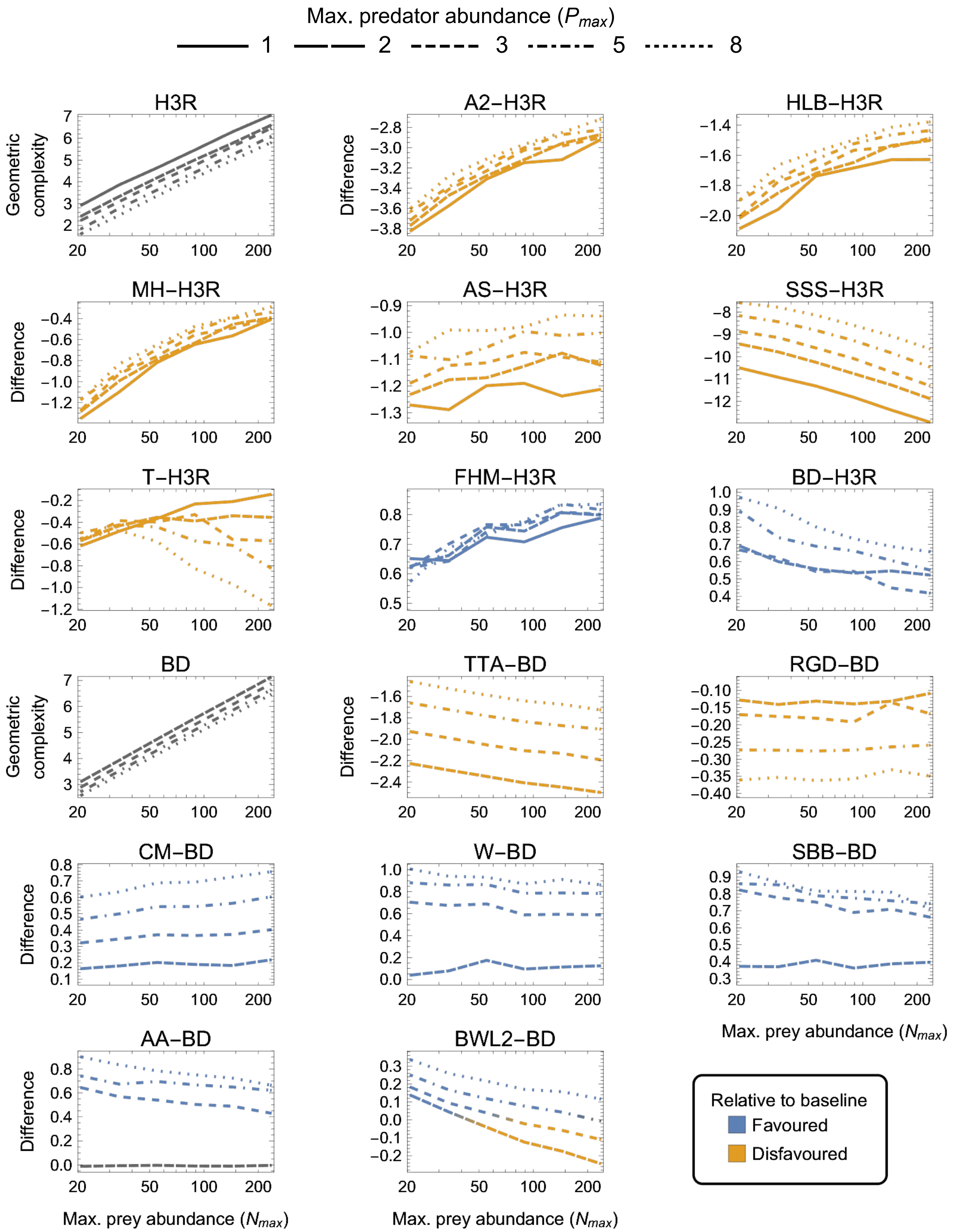

Figure S12: Absolute and relative geometric complexity for the three-parameter ($k = 3$) models with the constraint on the expected minimum number of eaten prey in the $N_{max}$ treatments being decreased by an order of magnitude (i.e. $\mathbb{E}[F(N_{max}, P, \theta)PT] \geq 1/10$).

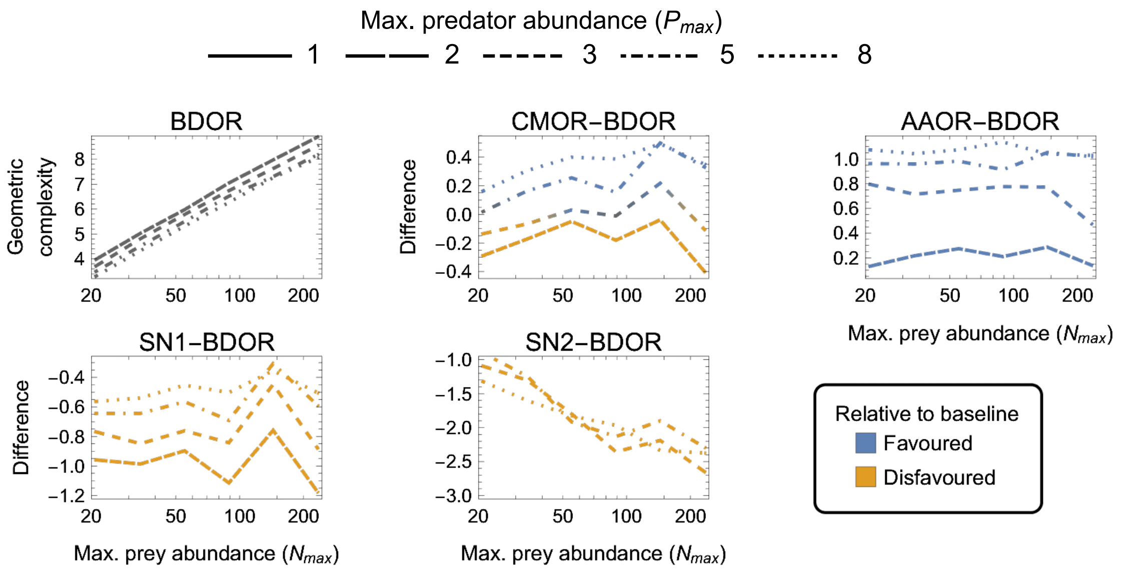

Figure S13: Absolute and relative geometric complexity for the four-parameter ($k = 4$) models with the constraint on the expected minimum number of eaten prey in the $N_{max}$ treatments being decreased by an order of magnitude (i.e. $\mathbb{E}[F(N_{max}, P, \theta)PT] \geq 1/10$).

### S8 Results for logarithmic designs with $\mathbb{E}[F(N,P,\theta)PT] \leq 10N_{max}$ constraint

Figures S14–S17 present the results for logarithmic experimental designs on which the constraint on the maximum expected number of eaten prey was increased by an order of magnitude from $\mathbb{E}[F(N,P,\theta)PT] \leq N_{max}$ to $\mathbb{E}[F(N,P,\theta)PT] \leq 10N_{max}$.

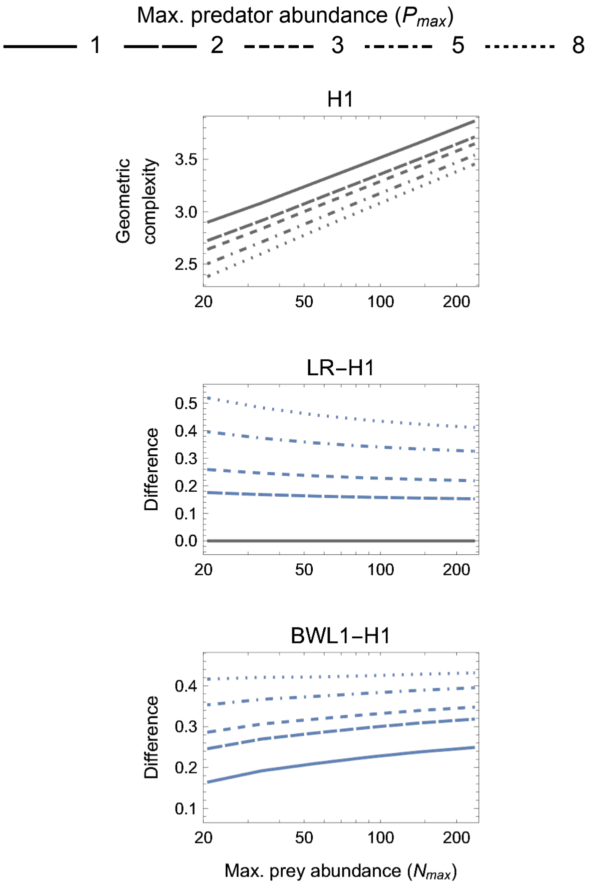

Figure S14: Absolute and relative geometric complexity for the one-parameter models ($k = 1$) with the constraint on the expected maximum number of eaten prey increased by an order of magnitude (i.e. $\mathbb{E}[F(N,P,\theta)PT] \leq 10N_{max}$).

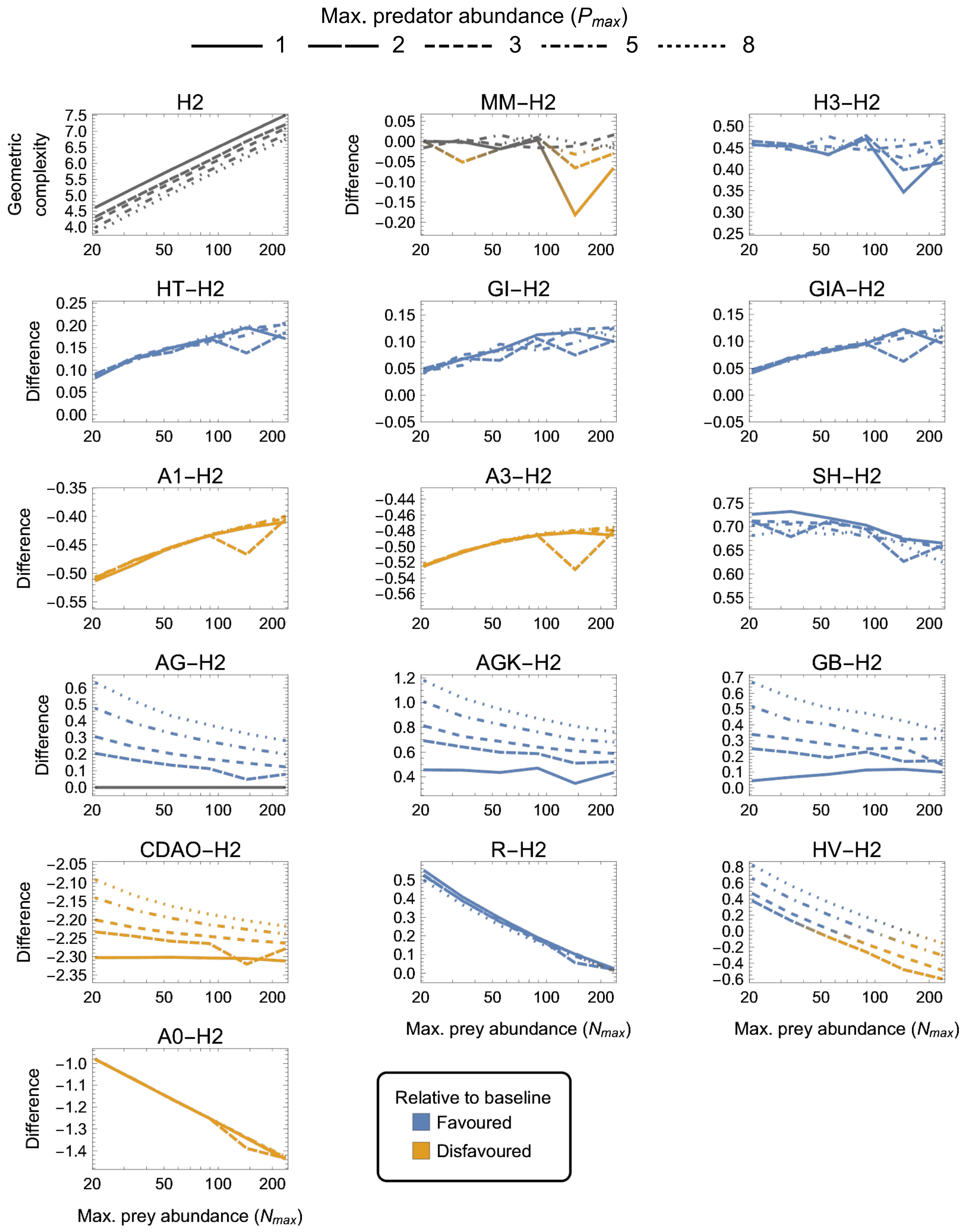

Figure S15: Absolute and relative geometric complexity for the two-parameter models ($k = 2$) with the constraint on the expected maximum number of eaten prey increased by an order of magnitude (i.e. $\mathbb{E}[F(N, P, \theta)PT] \leq 10N_{max}$).

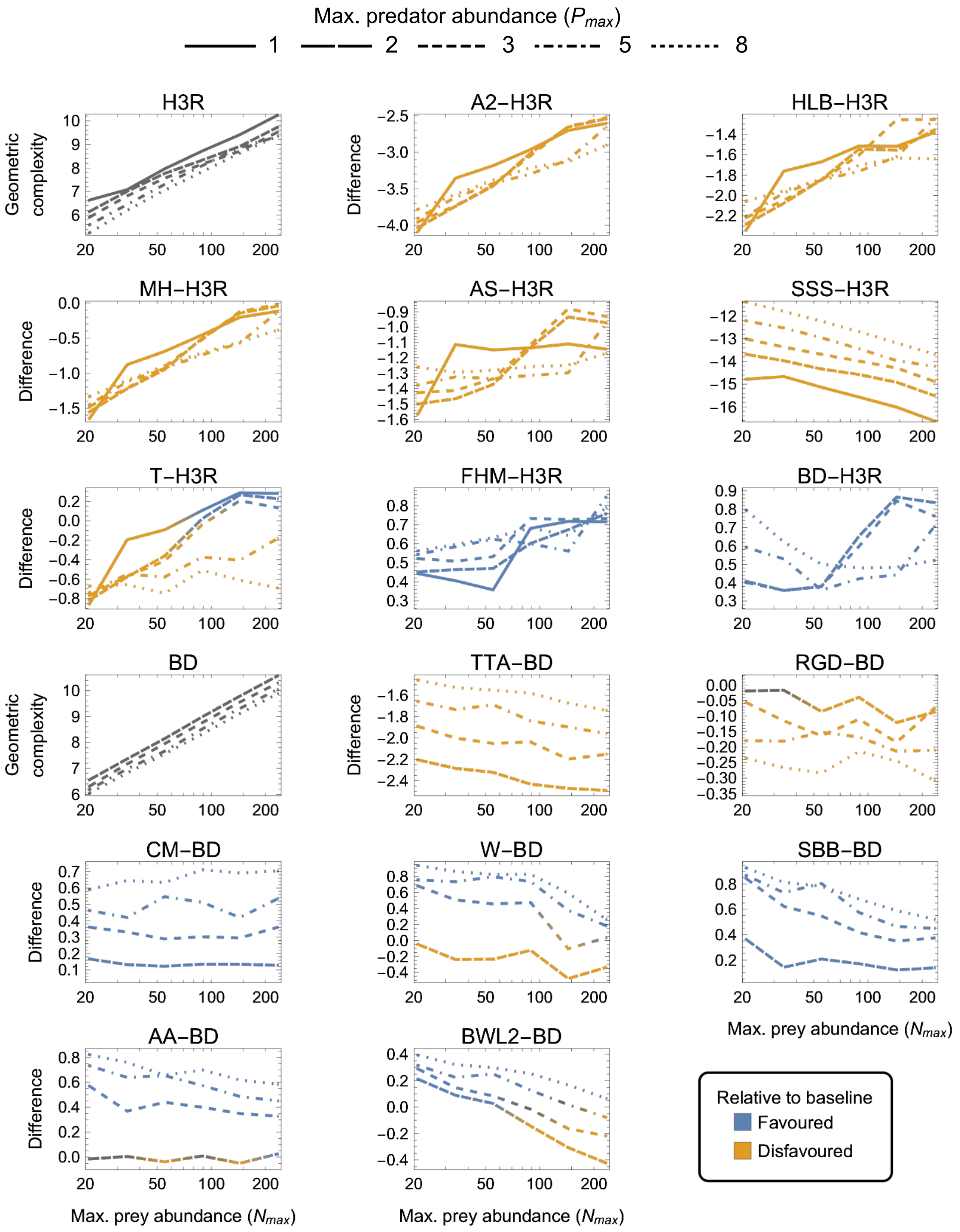

Figure S16: Absolute and relative geometric complexity for the three-parameter models ($k = 3$) with the constraint on the expected maximum number of eaten prey increased by an order of magnitude (i.e. $\mathbb{E}[F(N, P, \theta)PT] \leq 10N_{max}$).

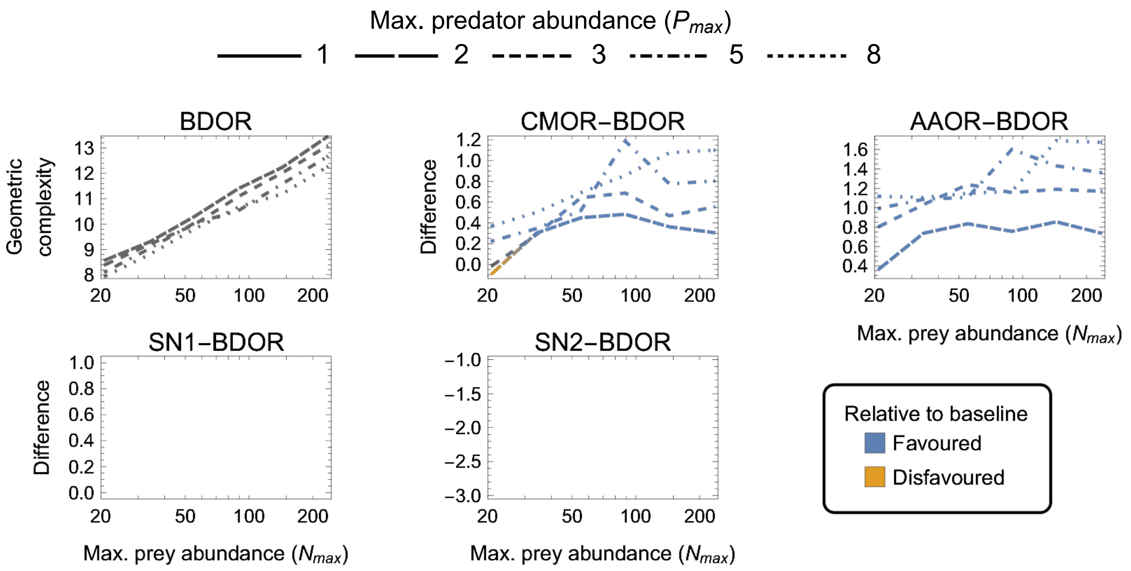

Figure S17: Absolute and relative geometric complexity for the four-parameter models ($k = 4$) with the constraint on the expected maximum number of eaten prey increased by an order of magnitude (i.e. $\mathbb{E}[F(N, P, \theta)PT] \leq 10N_{max}$).

### S9 Geometric complexity in non-replacement experiments

Our analyses of the main text only considered experimental designs in which eaten prey are continually replaced (or in which prey abundance is sufficiently well approximated by a constant value). Based on our own recent compilations of published functional-response data sets and consideration of replacement and non-replacement models, replacement designs are the more commonly used design and have more desirable statistical properties (Novak & Stouffer, 2021; Stouffer & Novak, 2021; as well as and citations therein). However, many studies do use non-replacement experimental designs.

Given that a binomial likelihood (appropriate for non-replacement studies) introduces additional non-linearity in the statistical model that is not present when using a Poisson likelihood (Novak & Stouffer, 2021), we expect non-replacement models to generally exhibit even greater variation in flexibility than observed in our main analysis. Unfortunately, most models applied to non-replacement scenarios do not have a closed-form solution and must be integrated numerically. Their likelihood function is therefore also not analytically accessible, making calculation of their FIA analytically intractable.

That said, a few functional-response models do have closed-form solutions in non-replacement scenarios. Of these, the most frequently assessed are the analogues to Holling's H2 and H3 replacement models: the Type II random predator equation of Rogers (1972) and its Type III form (Kratina *et al.*, 2009; Okuyama & Ruyle, 2011)[1]. These respectively describe the expected *proportion of initial prey* that are eaten as

$$p_{R2} = 1 - \frac{1}{ahN_0} W_0 \left( ahN_0 e^{-a(PT - hN_0)} \right) , \tag{S12}$$

where $W_0(x)$ represents the Lambert $W$ function and $N_0$ is the prey's abundance at the start of the experiment, and

$$p_{R3} = \frac{S - \sqrt{S^2 - 4a^2hN_0^3PT}}{2ahN_0^2} = \frac{2aN_0PT}{S + \sqrt{S^2 - 4a^2hN_0^3PT}} , \tag{S13}$$

where $S = 1 + aN_0PT + ahN_0^2$. Note that the first given form of Eqn. (S13) is the original form derived by Kratina *et al.* (2009) and Okuyama & Ruyle (2011) while the second form is an equivalent solution arrived at using the Citardauq formula (which we implemented).

A comparison of these two models for "variable" experimental designs (Fig. S18) indicates that Rogers' Type III is uniformly more flexible than Rogers' Type II (though less so than is H3 over H2), that their difference in flexibility increases with maximum prey abundances $N_{max}$ (in contrast to H2 and H3 whose difference is insensitive to $N_{max}$), and that their difference is insensitive to maximum prey abundance $P_{max}$ (consistent with H2 and H3). Their comparison for "fixed" experimental designs (Fig. S19) indicates that the greater flexibility of Rogers' Type III decreases with the number of prey treatment levels $L_N$ for low $P_N$, but is relatively insensitive to $L_N$ for large $P_N$ designs (in contrast to H2 and H3 whose difference slightly increases with $L_N$ and in insensitive to variation in $P_N$).

[1]Note that the Type III solution of Juliano (2001) inappropriately assumes that the number of prey remaining over time can be approximated by the number of prey at the start of the experiment (Rosenbaum & Rall, 2018).

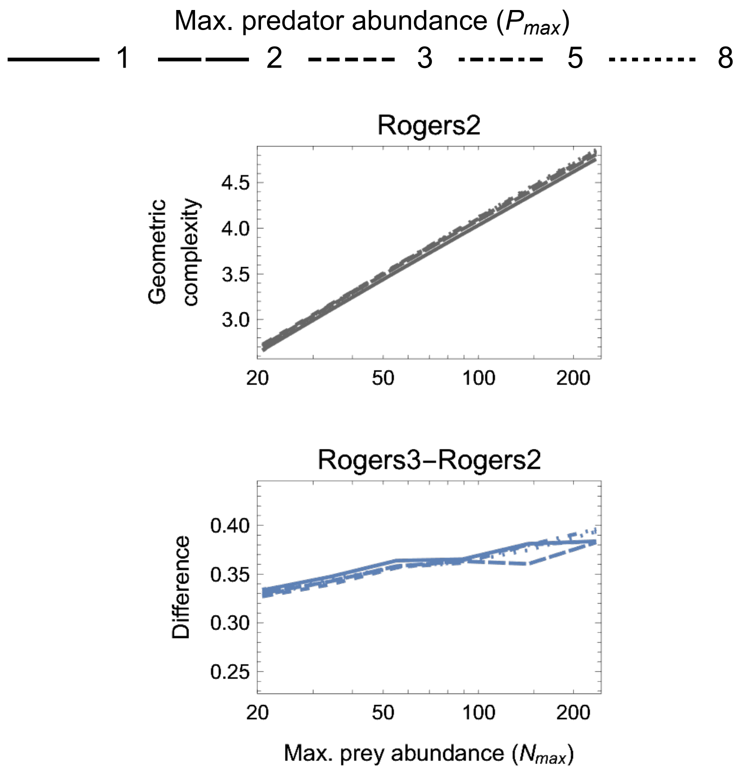

Figure S18: The geometric complexity of Rogers' random predator equation (the closed-form non-replacement analogue to the H2 model) and the relative geometric complexity of its Type III equivalent (the closed-form non-replacement analogue to the H3 model) for non-replacement experimental designs varying in the maximum number of (initial) prey and predator individuals ($N_{max}$ and $P_{max}$).

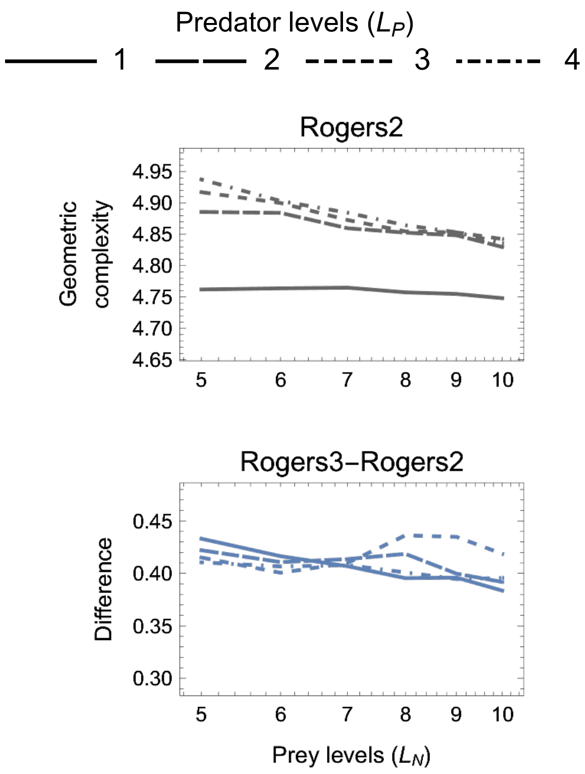

Figure S19: As in Fig. S18 but for experimental designs varying in the number of treatment levels ($L_N$ and $L_P$).

### References

Abrams, P. A. (1990). The effects of adaptive-behavior on the type-2 functional-response. *Ecology*, 71, 877–885.

Andrews, J. F. (1968). A mathematical model for the continuous culture of microorganisms utilizing inhibitory substrates. *Biotechnology and Bioengineering*, 10, 707–723.

Barbier, M., Wojcik, L. & Loreau, M. (2021). A macro-ecological approach to predation density-dependence. *Oikos*, 130, 553–570.

Beck, J. V. & Arnold, K. J. (1977). *Parameter Estimation in Engineering and Science.* John Wiley & Sons, Inc., New York, NY, USA.

Beddington, J. R. (1975). Mutual interference between parasites or predators and its effect on searching efficiency. *The Journal of Animal Ecology*, 44, 331–340.

Cole, D. (2020). *Parameter Redundancy and Identifiability.* CRC Press, Boca Raton, FL, USA.

DeAngelis, D. L., Goldstein, R. A. & O'Neill, R. V. (1975). A model for trophic interaction. *Ecology*, 56, 881–892.

Fujii, K., Holling, C. & Mace, P. (1986). A simple generalized model of attack by predators and parasites. *Ecological Research*, 1, 141–156.

Hassell, M. P., Lawton, J. H. & Beddington, J. R. (1977). Sigmoid functional responses by invertebrate predators and parasitoids. *The Journal of Animal Ecology*, 46, 249–262.

Jassby, A. D. & Platt, T. (1976). Mathematical formulation of the relationship between photosynthesis and light for phytoplankton. *Limnology and Oceanography*, 21, 540–547.

Jeschke, J. M., Kopp, M. & Tollrian, R. (2002). Predator functional responses: Discriminating between handling and digesting prey. *Ecological Monographs*, 72, 95–112.

Juliano, S. A. (2001). Nonlinear curve fitting: predation and functional response curves. *Design and analysis of ecological experiments*, 2, 178–196.

Kratina, P., Vos, M., Bateman, A. & Anholt, B. R. (2009). Functional responses modified by predator density. *Oecologia*, 159, 425–433.

Novak, M. & Stouffer, D. B. (2021). Systematic bias in studies of consumer functional responses. *Ecology Letters*, 24, 580–593.

Okuyama, T. & Ruyle, R. L. (2011). Solutions for functional response experiments. *Acta Oecologica*, 37, 512–516.

Pitt, M. A., Myung, I. J. & Zhang, S. (2002). Toward a method of selecting among computational models of cognition. *Psychological review*, 109, 472

Rissanen, J. J. (1996). Fisher information and stochastic complexity. *IEEE Transactions on Information Theory*, 42, 40–47.

Rogers, D. (1972). Random search and insect population models. *Journal of Animal Ecology*, 41, 369–383.

Rosenbaum, B. & Rall, B. C. (2018). Fitting functional responses: Direct parameter estimation by simulating differential equations. *Methods in Ecology and Evolution*, 9, 2076–2090.

Shapiro, A. (1986). Asymptotic theory of overparameterized structural models. *Journal of the American Statistical Association*, 81, 142–149.

Sokol, W. & Howell, J. A. (1981). Kinetics of phenol oxidation by washed cells. *Biotechnology and Bioengineering*, 23, 2039–2049.

Stouffer, D. B. & Novak, M. (2021). Hidden layers of density dependence in consumer feeding rates. *Ecology Letters*, 24, 520–532.

Tostowaryk, W. (1972). The effect of prey defense on the functional response of *Podisus modestus* (Hemiptera: Pentatomidae) to densities of the sawflies *Neodiprion swainei* and *N. pratti banksianae* (Hymenoptera: Neodiprionidae). *The Canadian Entomologist*, 104, 61–69.

Tyutyunov, Y., Titova, L. & Arditi, R. (2008). Predator interference emerging from trophotaxis in predator–prey systems: An individual-based approach. *Ecological Complexity*, 5, 48–58.
